# Complementary constraints in germ and immune cells shape evolution of gene regulation and phenotype

**DOI:** 10.64898/2026.03.27.714809

**Authors:** Kimberly N Griffin, Kira L Marshall, Gabriel A Russell, John Attanasio, Delaney B Farris, Haoming Yu, Eric Fagerberg, Nivedita R Iyer, Rebecca Lee, Kaelyn D Sumigray, Nikhil Joshi, Andrew Wang, Bluma J Lesch

**Affiliations:** Department of Genetics, Yale School of Medicine; New Haven, 06510, USA; Department of Internal Medicine (Rheumatology, Allergy, and Immunology), Yale School of Medicine; New Haven, 06510, USA; Department of Immunobiology, Yale School of Medicine; New Haven, 06510, USA; Department of Obstetrics, Gynecology & Reproductive Sciences, Yale School of Medicine; New Haven, 06510, USA; Yale Cancer Center, Yale School of Medicine; New Haven, 06510, USA

**Author notes:** equal contribution (Kimberly N Griffin, Kira L Marshall, Gabriel A Russell).

## Abstract

New mutations arise in germ cells and must preserve fertility while altering somatic phenotype to confer a fitness advantage. To understand how whole organisms balance these constraints, we generated mice with humanized noncoding promoter sequence at *Traf6*, an NF-κB pathway component with recently diverged chromatin state in mouse germ cells. The humanized allele (Traf6^h^) reinstated human-like expression in germ and immune cells, which we traced to a 13 base pair deletion that impairs CTCF binding. Traf6^h^ had no effect on fertility but induced a sensitized innate immune response to endotoxin. Other NF-κB pathway genes followed a similar pattern, suggesting that their expression has evolved in tandem and consistent with known mouse-human differences in inflammatory response. Our results reveal distinct roles for germ and immune cells in shaping gene regulatory evolution, where immune cells quickly adjust phenotype, enabling selection, while germ cells robustly maintain fertility despite changing expression levels, enabling inheritance.

## Introduction

Divergent gene regulation is a major source of evolutionary change ^1,2^. Regulatory divergence is caused by changes in noncoding sequence and enacted through assembly of alternative chromatin domains at regulatory elements, including enhancers and promoters ^3–10^. Importantly, evolutionary divergence in regulatory sequence impacts chromatin state and gene regulation differently in different cell types. While genomic sequence is the same in each cell of an organism, chromatin state induced at a given sequence differs from cell to cell, enabling diversity of cellular form and function ^11^. Therefore, selection for function in one tissue can unpredictably alter expression in other tissues, resulting in pleiotropic effects. This pleiotropy requires that multiple tissues coordinate phenotypic responses to the same change in regulatory sequence and complicates the relationship between evolution of regulatory sequence, chromatin state, and phenotype, making the outcomes of regulatory divergence difficult to predict ^12–15^.

The pleiotropy problem is especially acute for broadly expressed genes, which usually have important roles in multiple cell types. When expression of these genes is altered by an acquired regulatory mutation, the impact could be positive, negative, or neutral in different tissues, and the interplay of these effects ultimately determines the fitness of the organism and the likelihood that the mutation will persist. The necessity of balancing competing effects across tissues likely contributes to the longstanding observation that genes with broad expression patterns evolve more slowly than tissue-specific genes ^16–18^.

Among the many cell types in multicellular organisms, gametes (sperm and eggs) have a unique and central function in propagating the species and therefore represent a critical barrier in the evolution of broadly expressed genes. New mutations absolutely must be compatible with gamete development and fertility in order to persist. At the same time, male germ cells are transcriptionally promiscuous: spermatogenic cells express most of the genome at low levels, so mutations that impact expression of almost any gene have the potential to impair gamete function and fertility ^19–23^. How evolutionary change at broadly expressed genes occurs despite this germline barrier remains a fundamental question.

In mammalian gametes and gamete precursors, genes important for somatic development are marked by a specialized chromatin state called bivalency ^24–27^. Bivalency is defined by simultaneous enrichment of activation- and repression-associated histone modifications at the same site: H3 lysine 4 trimethylation (H3K4me3) and H3K27 trimethylation (H3K27me3), respectively ^24–26,28–30^. Bivalency in germ cells evolves in parallel with expression of the associated gene in somatic tissues, linking gamete chromatin state to regulatory divergence across different somatic tissues ^31,32^.

Here, we used the bivalent state in gametogenic cells as an entry point to ask how divergence in regulatory state occurs at broadly expressed genes in mammalian evolution. Genes that recently gained bivalency in gametes during evolution are commonly broadly expressed, while genes with deeply conserved bivalency are tissue-specific, suggesting that recent gain of bivalency marks broadly-expressed genes with the potential to evolve tissue-restricted expression. We selected *Traf6* (TNF receptor associated factor 6), a representative broadly-expressed gene that recently gained bivalency in murine germ cells, to interrogate the cross-tissue effects of regulatory sequence divergence. We generated humanized mice where the mouse *Traf6* promoter was replaced with orthologous human sequence and found that the humanized promoter sequence (Traf6^h^) was sufficient to convert both chromatin and gene expression to a human-like state in mouse testis. Despite strongly increased *Traf6* expression, mice homozygous for the Traf6^h^ promoter were normally fertile, indicating that the change in *Traf6* expression had minimal impact on germ cell function. However, elevated expression driven by *Traf6^h^* in immune cells led to significantly increased inflammatory phenotype. We traced the altered regulatory state to a 13 base pair deletion in the rodent lineage that eliminates a binding site for CCCTC-binding factor (CTCF). Genes involved in NF-κB and Toll-like receptor signaling followed a similar expression pattern to *Traf6*, suggesting that evolutionary changes in expression are coordinated across these functionally related genes and preferentially affect immune and germ cells. Our results provide a model for how potent somatic adaptations can evolve without imposing a penalty on fertility.

## Results

### The murine *Traf6* promoter acquired H3K27me3 to gain bivalency

We first sought to identify broadly-expressed genes that have undergone recent regulatory change during evolution. We mined our multi-species ChIP-seq dataset from male germline cells ^31^ for loci marked by bivalency in germ cells, and assessed tissue specificity using a previously reported metric (tau ^33,34^) and multi-tissue expression data from ENCODE ^35,36^. We found that in general, genes marked by bivalency are more tissue-specific than the genome average (**Figure 1A, Table S1**). Interestingly, this association with expression breadth was dependent on the evolutionary age of the bivalent state. Genes with deeply conserved bivalency were strongly tissue-specific (median tau 0.8607 and 0.8043 in mouse and human, respectively), while genes bivalent only in a single species were even more broadly expressed than the genome average (median tau 0.6352 and 0.6006 in mouse and human) (**Figure 1B**). Thus, acquisition of bivalency in germ cells marks evolutionarily recent regulatory change at genes with broad expression patterns in somatic tissue.

**Figure 1.**
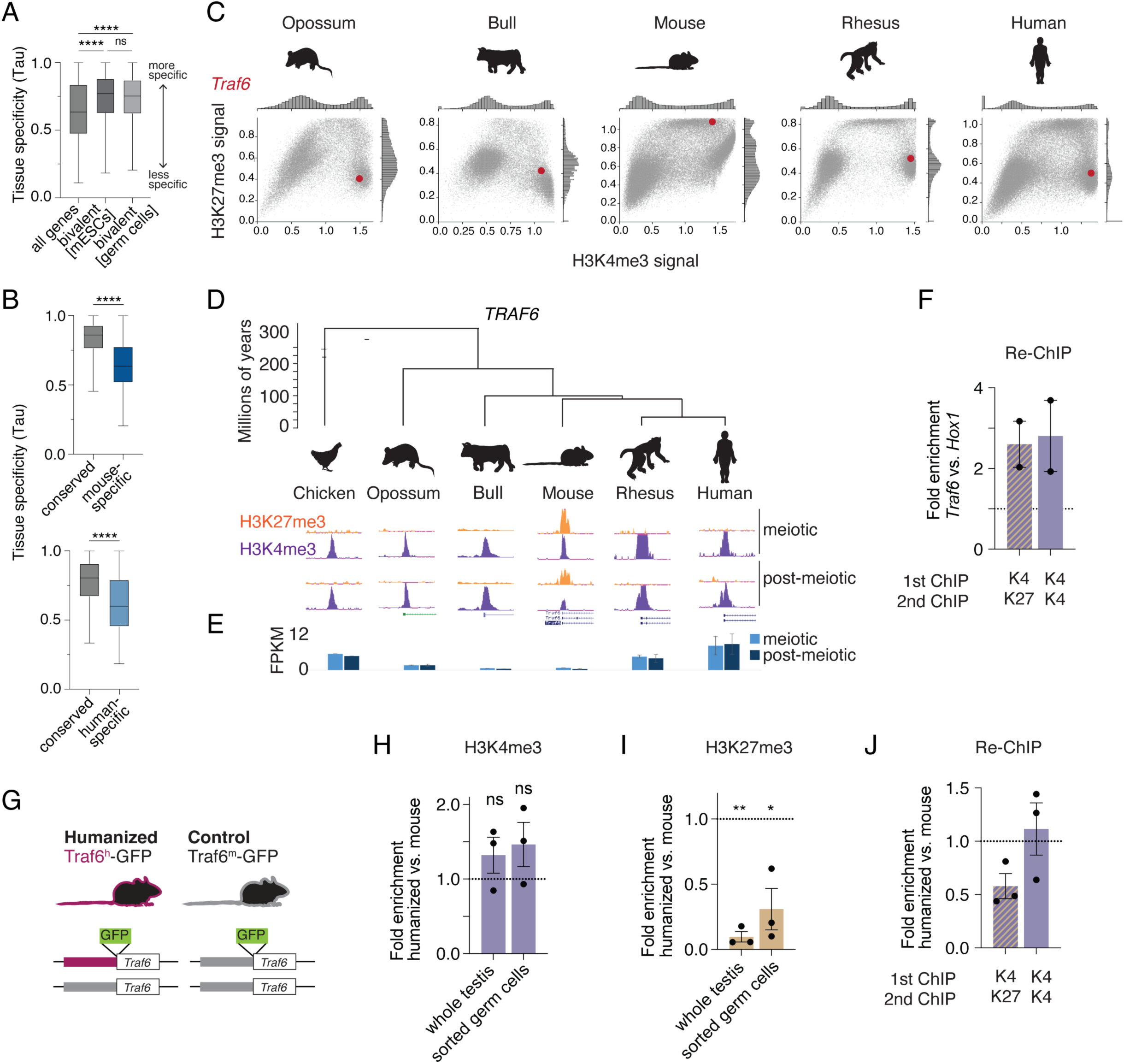
The Traf6 promoter gained H3K27me3 to become bivalent in the mouse lineage. **A**, Tissue specificity (tau) among all mouse genes or genes called as bivalent in mouse embryonic stem cells (mESCs) or germ cells. Higher tau corresponds to greater specificity. **B**, Tau values for mouse (top) or human (bottom) genes with conserved or species-specific bivalency in germ cells. For **A** and **B**, ****p<0.0001, unpaired t-test. **C**, ChIP-seq signal for H3K4me3 and H3K27me3 at all promoters in post-meiotic male germ cells of five mammalian species. *Traf6* promoter is shown in red. Data from GEO:GSE68507. **D**, Genome browser tracks at the *Traf6* promoter in meiotic and post-meiotic male germ cells in each species. **E**, Expression level (Fragments per kilobase million, FPKM) in male germ cells of each species. **F**, Re-ChIP for H3K4me3 followed by either H3K27me3 or a second H3K4me3 precipitation at the *Traf6* promoter. Values are normalized to a neutral locus (mm10 chr19:51758385-58758494) and plotted relative to a positive control for bivalency (*Hox1*). n=2 biological replicates. **G**, Cartoon showing humanization of the murine *Traf6* promoter at the endogenous locus. 1475bp of orthologous human sequence replaces the endogenous mouse promoter sequence on one allele. *Egfp* is inserted immediately downstream of the humanized (Traf6^h^-GFP) or endogenous mouse (Traf6^m^-GFP) sequence. The second allele is wild type. **H** and **I**, ChIP-qPCR for H3K4me3 (**H**) and H3K27me3 (**I**) at the humanized promoter relative to the mouse promoter in Traf6^h^-GFP mixed testis or sorted post-meiotic (round spermatid) germ cells. N=3 biological replicates. *p<0.05, **p<0.01, n.s. not significant at a threshold of p<0.05 by one-sample t-test relative to an expected value of 1. **J**, Re-ChIP at the humanized *Traf6* promoter in Traf6^h^-GFP testis cells. n=3 biological replicates. In **H-J**, data were normalized to a negative control locus (mm10 chr19:51758385-58758494) and the ratio of signal at the humanized vs. mouse allele is shown.

We next selected a representative locus to experimentally interrogate the factors that guide this regulatory evolution. We identified the *Traf6* promoter as broadly expressed (tau = 0.4768-0.6895) and enriched only for H3K4me3 in most mammalian species from primates to marsupials, but enriched for both H3K4me3 and H3K27me3 in mouse germ cells, indicating a switch from an active to a bivalent state in the murine lineage (**Figure 1C, Table S2**). The species difference in chromatin marks was constant across meiotic and post-meiotic development in male germ cells and persisted into epididymal sperm (**Figure 1D, S1A**). As expected for conversion from an active, H3K4me3-only to a bivalent state, mouse germ cells had low levels of *Traf6* transcript relative to other species (**Figure 1E**). We confirmed that the overlapping H3K4me3 and H3K27me3 ChIP-seq signals reflect true bivalency using sequential ChIP in mouse testis tissue (**Figure 1F**).

### Humanization of the mouse Traf6 promoter sequence recapitulates human regulatory state

To understand the mechanism and functional consequences of bivalency gain at *Traf6*, we generated a humanized allele (Traf6^h^-GFP) where 1171 base pairs of mouse *Traf6* promoter sequence underlying the H3K4me3 and H3K27me3 signal were replaced with the orthologous 1475 base pairs of human sequence (**Figure S1B-E**). We inserted an *Egfp* coding sequence immediately downstream of the humanized region to track expression, and left one allele of endogenous mouse *Traf6* intact for qPCR normalization (**Figure 1G**). We validated a change in chromatin state from mouse-like bivalent to human-like H3K4me3-only by ChIP-qPCR and sequential ChIP-qPCR for H3K4me3 and H3K27me3 in whole testis tissue and in sorted post-meiotic germ cells (round spermatids) of mice heterozygous for Traf6^h^-GFP. The results confirmed that H3K27me3 was depleted from the humanized compared to the mouse allele, while H3K4me3 enrichment was maintained at similar levels (**Figure 1H-J, S2A-B**).

The switch from bivalency to H3K4me3-only chromatin is predicted to increase gene expression. We therefore asked if the switch in chromatin state at the Traf6^h^ allele impacted expression of the GFP reporter. As a control for any effects of the *Egfp* sequence insertion, we generated a second allele, Traf6^m^-GFP, where the *Egfp* sequence is inserted in the same location in the absence of humanized sequence, putting it under control of the endogenous mouse *Traf6* promoter (**Figure 1E, S1C-D**). We confirmed that the promoter of the Traf6^m^-GFP allele was bivalent as expected (**Figure S3C**). GFP was highly expressed in the majority of testis cells carrying the Traf6^h^-GFP but not the Traf6^m^-GFP allele (**Figure 2A-B**). A similarly strong gain of GFP expression with Traf6^h^-GFP compared to Traf6^m^-GFP was detected by immunofluorescence in testis tissue (**Figure 2C**). At the transcript level, *Egfp* was the most upregulated transcript in Traf6^h^-GFP compared to Traf6^m^-GFP animals by RNA-seq (**Figure 2D, Table S3**). Notably, upregulation of the *Egfp* reporter was not matched by upregulation of the downstream *Traf6* transcript itself, confirming that our reporter lines were not confounded by downstream effects of *Traf6* overexpression (**Figure 2E**).

**Figure 2.**
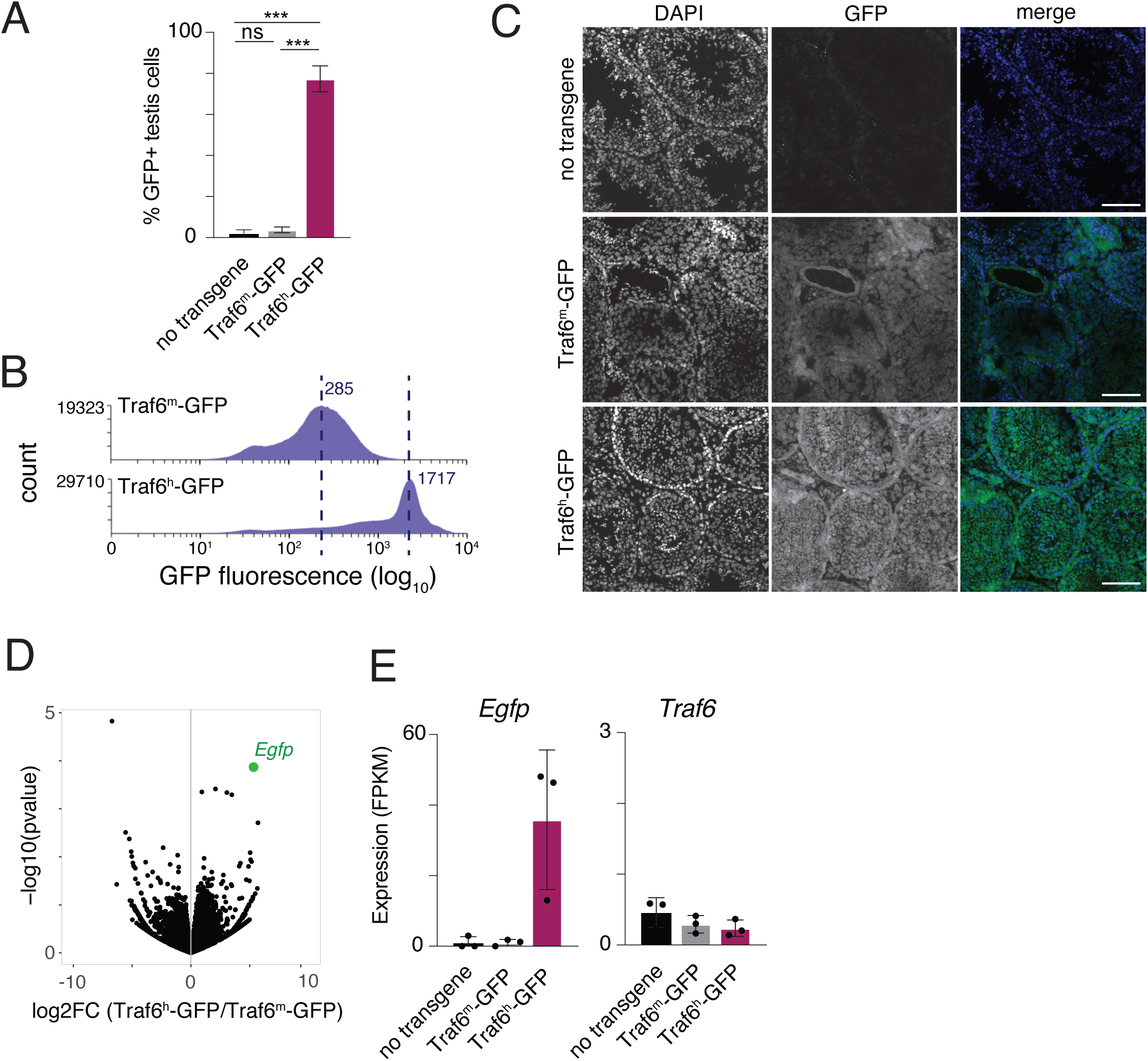
The humanized Traf6 promoter drives increased expression. **A**, Percentage of cells expressing GFP in wild type, Traf6^m^-GFP and Traf6^h^-GFP testis by flow cytometry. ***p<0.001, Welch’s t-test. **B**, Flow cytometry histograms for GFP in dissociated testis cells from Traf6^m^-GFP and Traf6^h^-GFP mice. Dashed line indicates median signal. **C**, Immunofluorescence staining for GFP in fixed testis tissue from wild type, Traf6^m^-GFP and Traf6^h^-GFP mice. DNA is counterstained with DAPI. Scale bar, 100um. **D**, Volcano plot from RNA-seq data from testis tissue of Traf6^h^-GFP compared to Traf6^m^-GFP mice. Each point represents a transcript; *Egfp* transcript is highlighted. **E**, Quantitation of RNA-seq transcript levels for *Egfp* and *Traf6*. n=3 biological replicates.

### Humanized *Traf6* regulation supports normal spermatogenic development and fertility

Having validated that the humanized Traf6^h^ promoter allele recapitulates a human-like chromatin and expression state in germ cells, we next asked how the humanized *Traf6* promoter impacts phenotype. Since our GFP insertion interferes with regulation of the *Traf6* gene (**Figure 2E**), we generated a Traf6^h^ allele with a humanized promoter sequence but no *Egfp* insertion, and homozygosed this allele (Traf6^h/h^) to better study phenotype (**Figure 3A**). Traf6^h/h^ mice were viable and healthy with no apparent morphological or physiologic abnormalities in either sex.

**Figure 3.**
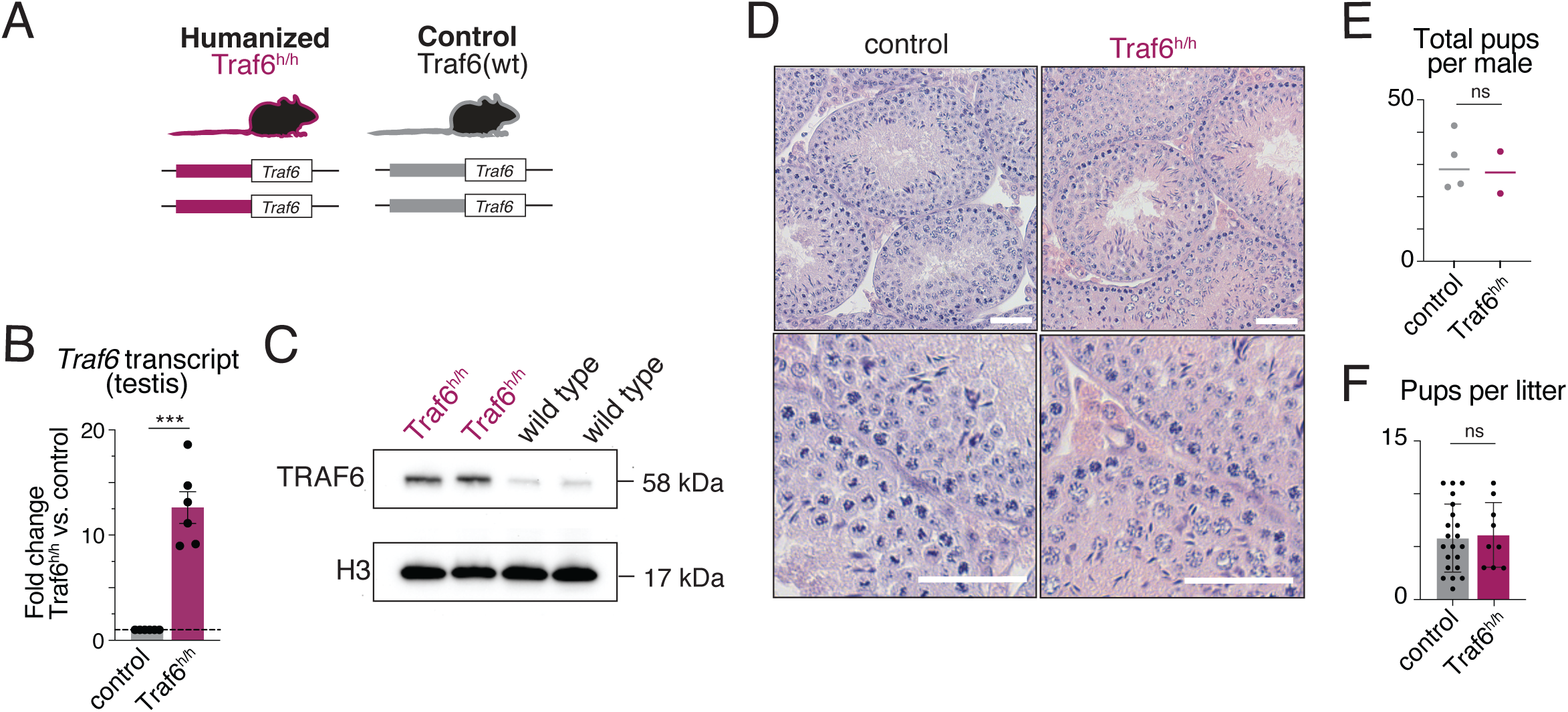
Traf6h/h mice have normal fertility. **A**, Schematic of Traf6h/h homozygous humanized and control (wild type) Traf6 promoter genotypes. B, Relative expression (RT-qPCR) of Traf6 transcript in testis tissue of Traf6h/h homozygous compared to control animals. n=6 biological replicates. ***p<0.001, 1-sample t-test. Error bars represent s.e.m.; dashed line is at y=1 (no change). Actb was used as an internal control. C, Western blot for TRAF6 in Traf6h/h and wild type testes. n=2 biological replicates for each genotype are shown. H3 is a loading control. D, H&E staining of testis sections from Traf6h/h and control animals. All spermatogenic cell types are present and flagellated sperm is visible in the lumen of the seminiferous tubules. Scale bar, 50um. E and F, Total pups produced per male (E) and pups per litter (F) after 6 months of continuous mating to wild type females. ns, not significant at p<0.05 by unpaired t-test.

We first assessed the impact of Traf6^h^ on the primary function of the germline, fertility, under standard laboratory conditions. We confirmed that *Traf6* transcript and protein expression are elevated in Traf6^h/h^ testis tissue, reflecting the GFP expression we observed with the reporter allele (**Figure 3B-C**). Histological examination of Traf6^h/h^ testis tissue revealed normal spermatogenesis, with all spermatogenic cell types present in the seminiferous epithelium and normal-appearing sperm in the tubule lumen (**Figure 3D**). Consistent with normal spermatogenesis, there was no detectable defect in fertility as Traf6^h/h^ male mice generated normal numbers of pups (**Figure 3E-F**), and we observed that fertility of female Traf6^h/h^ breeders was likewise unremarkable. We conclude that humanization of the *Traf6* promoter robustly humanizes regulatory state and activates expression in testis, but has no biologically significant effect on testicular phenotype or fertility.

### Humanized *Traf6* regulation alters immune expression and inflammatory phenotypes

Since there was no strong effect on germ cell phenotype or fertility, we predicted that any functional effects of *Traf6* regulatory divergence would be found in somatic tissue. TRAF6 is an E3 ubiquitin ligase that is broadly expressed across tissues in mice and humans and has a range of functions, most prominently as a positive mediator of canonical NF-κB signaling ^37–40^. It is important for development and function of both adaptive and innate immune cells as well as organogenesis of the thymus and lymph node ^37,39–45^. To determine if humanized *Traf6* regulation alters immune function, we focused on the well-characterized role of TRAF6 as a regulator of Toll-like receptor 4 (TLR4) signaling in myeloid cells to enact a systemic inflammatory response (**Figure 4A**). Upon activation of TLR4, TRAF6 serves as an essential adaptor protein, coordinating the nuclear translocation of NF-κB and leading to the production of pro-inflammatory cytokines including TNFα and IL-6 ^46–50^. To examine how humanized regulation of *Traf6* impacts this process, we utilized lipopolysaccharide (LPS, endotoxin). LPS is a well-established ligand of TLR4, and LPS challenge is a commonly used mouse model of sepsis, wherein mortality results entirely from the ensuing systemic inflammatory response. Upon intraperitoneal challenge with LPS, Traf6^h/h^ mice were found to exhibit a dramatic increase in endotoxin sensitivity, resulting in complete (100%) mortality in response to the same LD50 dose given to controls (**Figure 4B**). Furthermore, Traf6^h/h^ mice had significantly elevated levels of circulating pro-inflammatory cytokines IL-6 and TNFα throughout challenge (**Figure 4C-D**), consistent with the role of TRAF6 as a positive regulator in the production of these cytokines, as well as the role of systemic inflammation as the cause of death in this model. Thus, humanization of *Traf6* regulation results in heightened responsiveness and mortality during systemic pro-inflammatory challenge.

**Figure 4.**
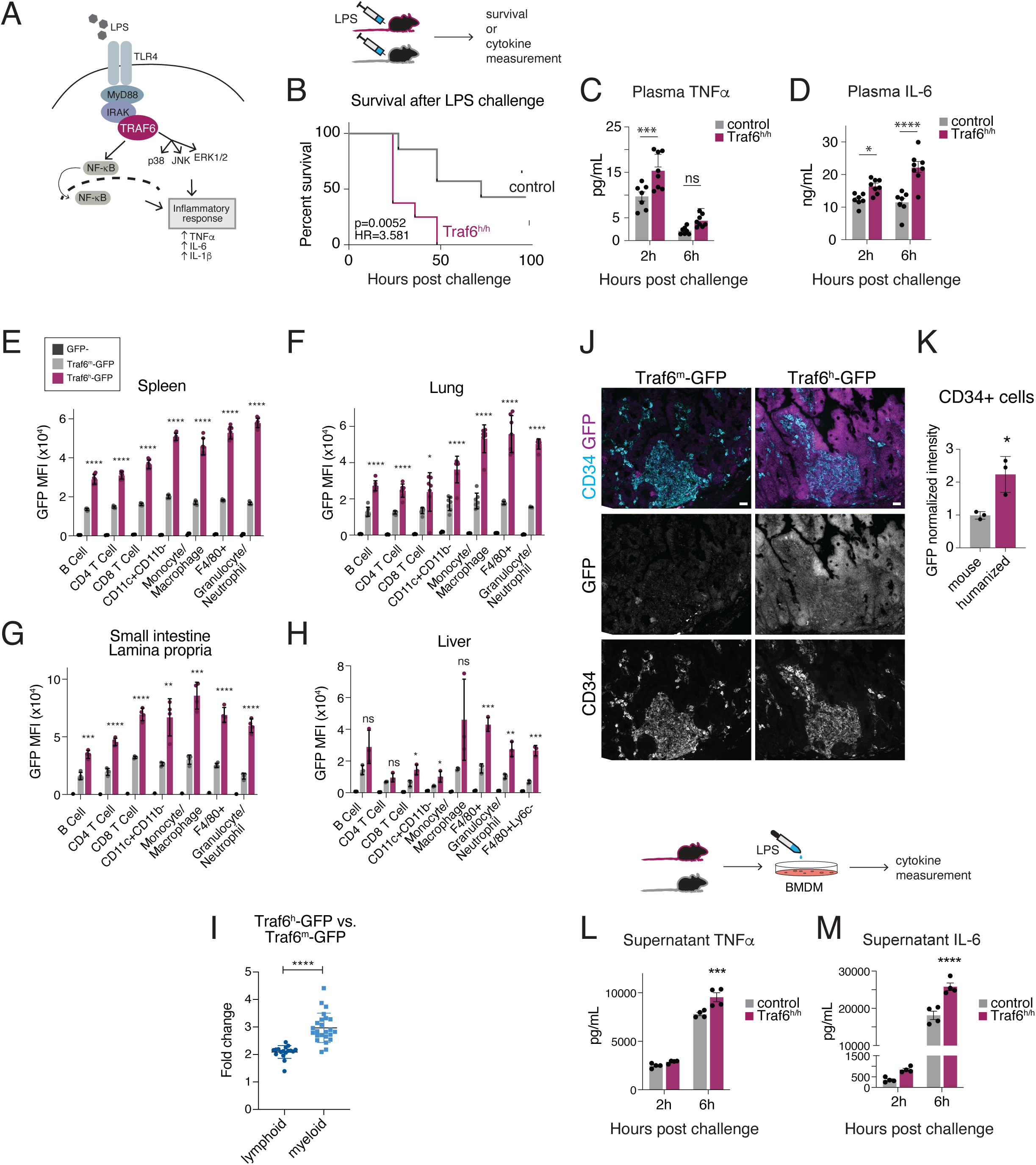
Traf6^h/h^ mice have a sensitized systemic inflammatory response to endotoxin. **A**, Cartoon of TRAF6 function in the NF-kB pathway. **B**, Survival of control or Traf6^h/h^ adult mice following intraperitoneal challenge with LD50 dose of LPS. n=7-8 over two independent trials. P-value calculated by Mantel-Cox test; HR = log-rank hazard ratio. A schematic of the experiment in B-D is shown above. **C** and **D**, Plasma TNFα (**C**) or IL-6 (**D)** levels at 2 or 6 hours after LPS challenge. n=7-8 over two independent trials. *p<0.05, ***p<0.001, ****p<0.0001, ANOVA with Sidak’s multiple comparisons test. **E** to **H**, Mean GFP fluorescence intensity for different populations of immune cells in spleen (**E**), lung (**F**), small intestine lamina propria (**G**), or liver (**H**) from Traf6^h^-GFP or Traf6^m^-GFP mice. *p<0.05, **p<0.01, ***p<0.001, ****p<0.0001, unpaired Welch’s t-test for Traf6^h^-GFP vs. Traf6^m^-GFP. MFI, mean fluorescence intensity. CD11c+CD11b-population indicates antigen-presenting cells (APC); F4/80+ population indicates differentiated macrophages. Gating strategy is shown in **Figure S3**. **I**, Ratios of GFP fluorescence intensity in Traf6^h^-GFP vs. Traf6^m^-GFP cells in lymphoid or myeloid populations across all tissues. Each point represents the ratio of one immune population in a single tissue. ****p<0.0001, unpaired Welch’s t-test. **J**, Representative images of small intestine epithelium immunostained for GFP (magenta) and the pan-immune marker CD34 (cyan) in Traf6^h^-GFP and Traf6^m^-GFP mice. Scale bar, 20 um. **K**, Mean GFP signal in Traf6^h^-GFP CD34+ (immune) cells, normalized to average Traf6^m^-GFP signal across n=3 biological replicates. *p<0.05, unpaired t-test. **L** and **M**, Supernatant TNFα (**L**) or IL-6 (**M)** levels at 2 or 6 hours after treatment of control or Traf6^h/h^ bone marrow derived macrophages (BMDMs) with 1ng/mL LPS. A schematic of the experiment is shown above. n=4 derived from n=2 biological replicates per genotype; data representative of two independent experiments. *p<0.05, ***p<0.001, ****p<0.0001, ANOVA with Sidak’s multiple comparisons test.

Sensitization to LPS challenge could be due to increased TRAF6 expression in effector cells, changes in relative proportions of different immune cell types, or both. To test for immune lineage bias, we first evaluated immune populations in several tissues and organs. We found no or minimal change in proportions of immune cell types in either myeloid or lymphoid lineages in spleen, bone marrow, lung, or liver tissue in Traf6^h/h^ mice compared to wild type controls (**Figure S3, S4A-D**).

We next asked if levels of *Traf6* expression were elevated within immune cells. We dissociated cells from liver, spleen, lung, intestine, and bone marrow isolated from mice carrying the Traf6^h^-GFP and Traf6^m^-GFP reporter alleles and assessed GFP expression in populations of circulating and tissue-resident immune cells by flow cytometry. Expression of GFP under the humanized promoter allele was higher than under the endogenous mouse promoter in nearly all immune populations (**Figure 4E-H, S4E-F)**. Reporter expression was increased in both myeloid and lymphoid lineages but the effect was consistently stronger in myeloid populations (fold change over Traf6^m^: 1.39-2.45 lymphoid, 2.09-4.41 myeloid, **Figure 4I**). We further confirmed elevated expression of Traf6^h^-GFP compared to Traf6^m^-GFP by immunofluorescence in small intestine, where GFP expression was significantly higher in immune as well as epithelial cells in Traf6^h^-GFP animals (**Figure 4J-K, S4G-H**). Together, these results indicate that the Traf6^h^ promoter regulates immune phenotypes by increasing expression of *Traf6* within immune cells rather than altering composition of the immune compartment.

We next asked if the increased expression driven by the Traf6^h^ promoter was largely selective for immune cells or if it reflected a nonspecific increase in promoter activity driving global upregulation. We assessed *Traf6* transcript levels in multiple somatic tissues and found that substantially increased levels of *Traf6* transcript were detected only in testis (8-19x relative to control, **Figure S4I**). Mild but statistically significant increases in transcript levels were also detected in lung, heart, and small intestine, consistent with modestly increased GFP expression seen in intestinal epithelial cells in addition to immune cells in our immunofluorescence data (**Figure S4G-H**). In lung and heart, this result could also indicate selective upregulation of *Traf6* in immune populations that is partially masked by non-immune cells at the bulk tissue level. We conclude that the humanized *Traf6* promoter drives elevated expression primarily in immune and germ cells.

Given the role of myeloid cells as first responders to immune stimulus and the finding that the Traf6^h^ promoter drives expression more strongly in myeloid populations, we then used a common in vitro model to functionally validate a stronger myeloid response to stimulus in Traf6^h/h^ immune cells. We generated bone marrow derived macrophages (BMDMs) from Traf6^h/h^ animals and littermate controls and challenged these cells with LPS in vitro. In response to LPS stimulation, Traf6^h/h^ BMDMs produced higher levels of pro-inflammatory cytokines IL-6 and TNFα relative to controls (**Figure 4L-M**), further confirming that the elevated response to LPS in Traf6^h/h^ mice is mediated by increased *Traf6* expression in immune cells.

### Loss of CTCF binding permits gain of H3K27me3 at the *Traf6* promoter

We next sought to determine how sequence divergence at the *Traf6* promoter mediates altered expression and chromatin state. We interrogated the 1475 bp sequence constituting the humanized promoter transgene for features that differentiate it from the orthologous sequence in mouse. GC content is known to contribute to recruitment of Polycomb and resultant enrichment of H3K27me3 in mouse ESCs (mESCs) ^51–53^, but the GC content of the two sequences is similar (human: 60.3%; mouse: 59.4%). Additionally, while transposable element insertions are a common source of regulatory change in evolution ^54–58^, we did not detect transposable element sequences in either the human or mouse *Traf6* promoter sequences. We noted that a predicted binding site for CTCF in primate *Traf6* promoter sequences (human, chimpanzee, and rhesus macaque) overlapped a natural deletion in the mouse genome (**Figure 5A**-**C**). Indeed, we found that the Traf6^h^ sequence recruits CTCF more strongly than the endogenous mouse sequence in testis at the predicted CTCF site using allele-specific ChIP-qPCR in our internally controlled heterozygous Traf6^h^-GFP mouse line (**Figure 5D**).

**Figure 5.**
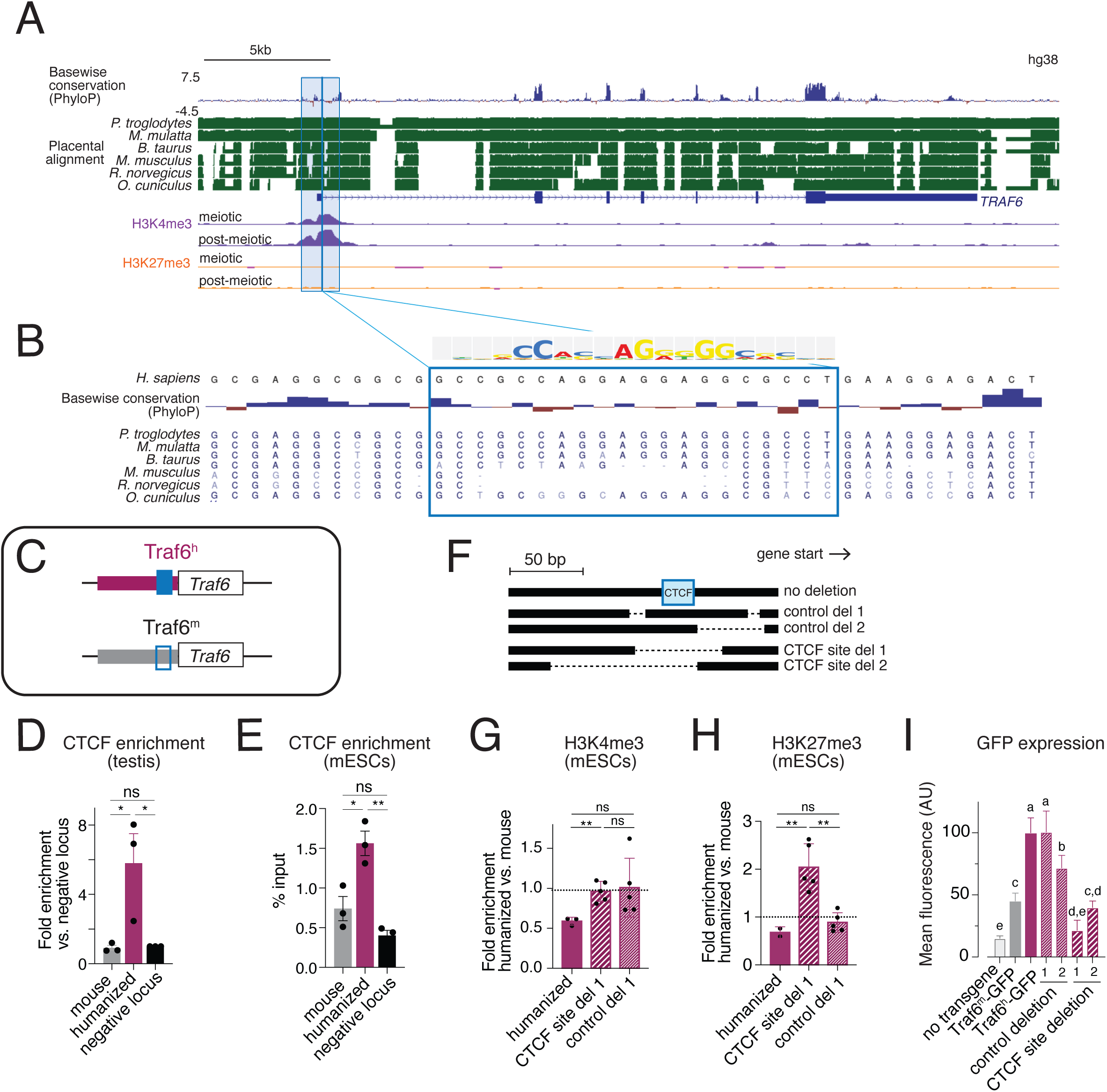
Loss of a putative CTCF binding site permits H3K27me3 deposition at the murine *Traf6* promoter. **A**, Sequence conservation scores between human and chimpanzee, rhesus macaque, bull, mouse, rat, and rabbit at the *Traf6* locus. H3K4me3 and H3K27me3 ChIP-seq signal from human meiotic and postmeiotic male germ cells is shown below. Box shows regions replaced by orthologous human sequence. **B**, CACTUS alignment ^109^ in the vicinity of a CTCF binding site (light blue box) present in primate but not mouse or rat sequence. TRANSFAC position weight matrix for the predicted CTCF binding site is shown above the alignment. **C**, Cartoon of CTCF site present in humanized allele and absent in mouse *Traf6* allele. **D**, CTCF ChIP-qPCR at the mouse and humanized Traf6 promoter sequences in Traf6^h^-GFP testis tissue. *p<0.05, ratio paired t-test, n=3 biological replicates. Data were normalized to a negative control locus (mm10 chr19:51758385-58758494) and the ratio of signal at the humanized vs. mouse allele is shown. **E**, CTCF ChIP-qPCR at the predicted CTCF binding site in mouse and humanized Traf6 promoter sequences in Traf6^h^-GFP mESCs. *p<0.05, **p<0.01, unpaired t-test, n=3 biological replicates. **F**, Cartoon of CRISPR deletions in the Traf6^h^-GFP allele relative to the putative CTCF binding site. **G** and **H**, H3K4me3 and H3K27me3 ChIP-qPCR at the humanized *Traf6* promoter, CTCF site deletion, and control site deletion. **p<0.01, Welch’s t-test. ns, not significant at p<0.05. n=3-5 biological replicates; data were normalized to a negative control locus (mm10 chr19:51758385-58758494) and the ratio of signal at the humanized vs. mouse allele is shown. **I**, Mean GFP fluorescence assayed by flow cytometry for wild type mESCs (no transgene), or mESCs heterozygous for the Traf6^m^-GFP or Traf6^h^-GFP alleles. p<0.0001, one-way ANOVA. Letters a-e indicate significance groups: conditions marked by the same letter are not significantly different from each other and all other comparisons are significant at p<0.05 by Tukey’s multiple comparisons test.

CTCF plays many roles in genome regulation, including mediating three-dimensional contacts and promotion of transcriptional activity ^59^. Inspection of micro-C data from mouse and human ESCs did not reveal meaningful species differences in local chromatin contacts near *Traf6*, indicating that changes in 3D contacts likely do not explain the observed differences in gene regulation (**Figure S5**). CTCF has also been found to directly antagonize H3K27me3 deposition, suggesting a mechanism where loss of CTCF binding permits gain of H3K27me3 deposition at the *Traf6* promoter ^60,61^. Consistent with an antagonistic relationship, genome-wide CTCF binding sites are enriched for H3K4me3 but depleted for H3K27me3 in both mouse and human testis (**Figure S6A-B**).

Next, we confirmed that the Traf6^h^ promoter is more strongly enriched for CTCF than the Traf6^m^ promoter in mESCs as in testis, using primers targeting either the site of the predicted CTCF motif or a nearby ENCODE-annotated CTCF peak (**Figure 5E, S6C-D**). To test if CTCF directly antagonizes H3K27me3 at the *Traf6* promoter, we generated mESCs carrying one copy of the Traf6^h^-GFP allele and used CRISPR to delete either the CTCF binding site or adjacent control sites within the Traf6^h^ sequence (**Figure 5F**). Deletion of the CTCF binding site, but not adjacent deletions that leave the CTCF site intact, increased H3K27me3 enrichment on the Traf6^h^ promoter sequence without changing H3K4me3 (**Figure 5G-H**). Consistent with this result, deletion of the CTCF binding site, but not adjacent control deletions, reduced Traf6^h^-GFP expression to levels similar to Traf6^m^-GFP (**Figure 5I**). These results suggest that the natural deletion compromising the CTCF binding site in the mouse *Traf6* promoter facilitates gain of H3K27me3 and reduces expression of *Traf6*.

### A preexisting H3K27me3 domain expanded into the *Traf6* promoter following loss of CTCF binding in the rodent lineage

Finally, we asked if the CTCF-positive, H3K27me3-negative configuration represents the ancestral state of the *Traf6* promoter, suggested by the absence of H3K27me3 at *Traf6* in other mammalian species (**Figure 1C-D**). To trace the evolutionary history at this locus, we collected H3K4me3, H3K27me3, and CTCF ChIP data from testis tissue of an additional rodent species, rat (*Rattus norvegicus*, 11.6 million years diverged from mouse) as well as rabbit (*Oryctolagus cuniculus*, 79 million years diverged from mouse and rat) (**Figure 6A, Table S4, Table S5**) ^62^. The rat *Traf6* promoter sequence resembles the mouse and carries the same CTCF binding site deletion. In the rabbit genome, the predicted CTCF site is intact but diverges from primate at several individual base pairs, making predictions of CTCF binding based on sequence in the rabbit ambiguous (**Figure 5B**). We interpret these sequence differences as resulting from a deletion that occurred in the rodent common ancestor after its split from the rabbit lineage, although we note that further analysis would be needed to conclusively determine the ancestral state.

**Figure 6.**
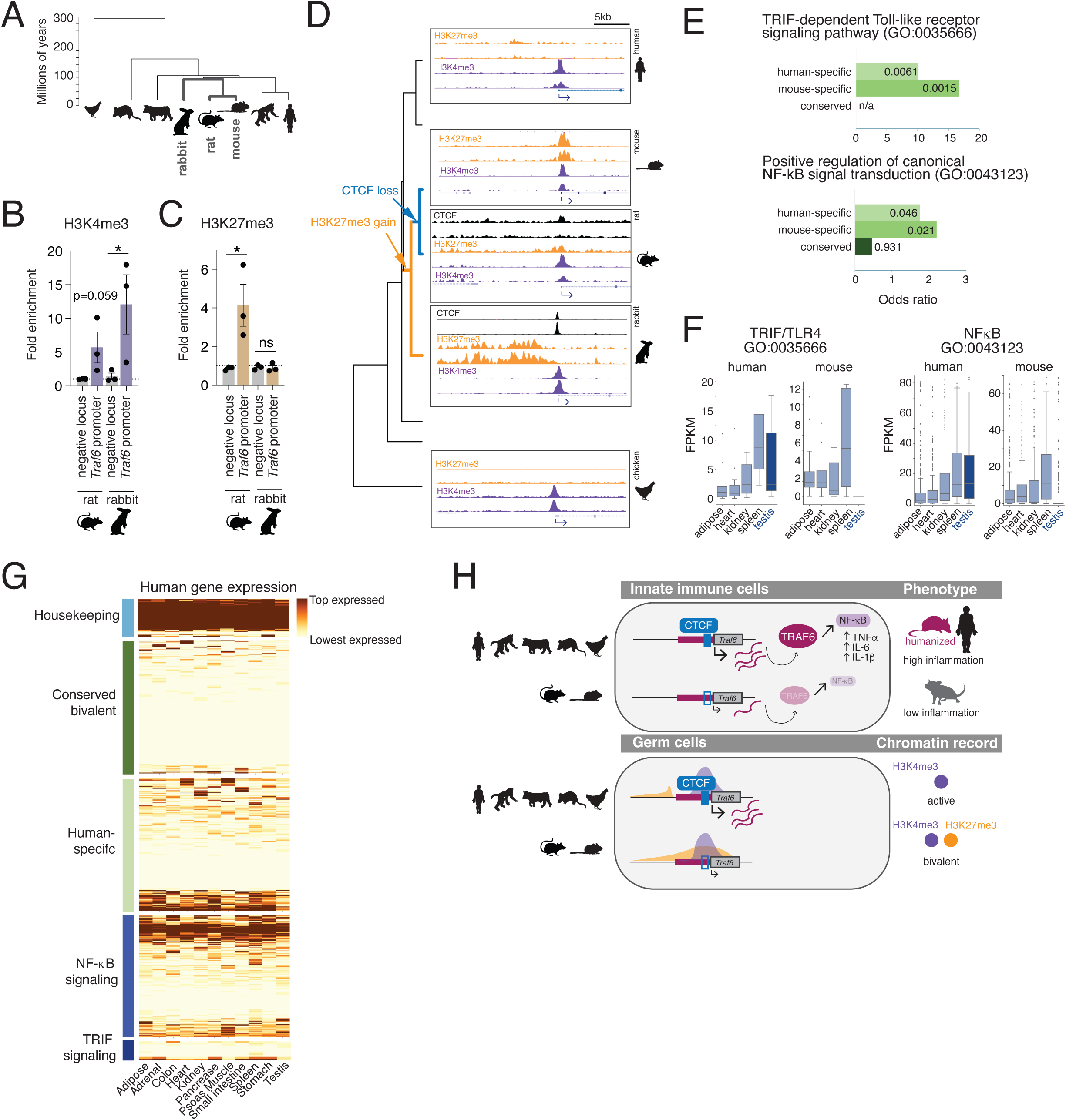
Evolutionary dynamics of gene regulation at *Traf6* and NF-kB pathway genes. **A**, Mammalian phylogeny showing relationship between rabbit, mouse, and rat, and inferred loss of CTCF site. **B** and **C**, H3K4me3 and H3K27me3 ChIP-qPCR at the *Traf6* promoter and a negative control locus (rat: rn6 chr1:272076197-272076337; rabbit: oryCun2 chr14:101335505-101335641) in rat and rabbit. *p<0.05, ratio paired t-test. ns, not significant at p<0.05. n=3 biological replicates; data represent fold enrichment relative to a second negative locus (rat: rn6 chr1:272081766-272082001; rabbit: oryCun2 chr1:61991839-61991978). **D**, ChIP-seq tracks for H3K4me3, H3K27me3, and CTCF at the *Traf6* locus in rat and rabbit. H3K4me3 and H3K27me3 data from human, mouse, and a chicken outgroup (*Gallus gallus*) are shown for comparison. Tracks show data from enriched germ cells except data from mixed testis cells for rat CTCF and rabbit. **E**, Enrichment for the GO categories “Positive regulation of canonical NF-κB signal transduction” (GO:0043123) and “TRIF-dependent Toll-like receptor signaling pathway” (GO:0035666) among genes with conserved, human-specific, or mouse-specific germline bivalency. p-values for enrichment are shown inside the bars (hypergeometric test). **F**, Expression (FPKM) of genes in the NF-κB and TRIF/TLR pathways from (**E**) in representative mouse and human tissues including spleen and testis (data from ENCODE, see **Table S1**). Boxes show interquartile range (IQR), whiskers extend to 1.5 IQRs from upper and lower quartile, and data beyond this range is plotted as points. **G**, Heatmap of gene expression decile across human tissues for housekeeping (n=100), conserved bivalent (n=349), human-specific bivalent (n=300, random sample out of 796) genes, and for genes in the GO categories “Positive regulation of canonical NF-kB signal transduction” (GO:0043123, n=322) and “TRIF-dependent Toll-like receptor signaling pathway” (GO:0035666, n=11). **H**, Proposed model for evolutionary changes to *Traf6* regulation in immune and germ tissues. Top, loss of CTCF binding in the rodent lineage reduced expression of TRAF6, NF-κB signaling, and inflammatory response in immune cells, providing a fitness advantage. Bottom, loss of CTCF binding permitted expansion of a nascent H3K27me3 domain in germ cells, generating a bivalent state that parallels the change in somatic gene function.

By ChIP-qPCR, both rat and rabbit promoters were enriched for H3K4me3, while only rat was enriched for H3K27me3 (**Figure 6B-C**). At the same time, the rabbit but not the rat promoter was enriched for CTCF (**Figure 6D**), supporting a history where CTCF binding was lost and H3K27me3 enrichment gained after the split between rabbits and true rodents. Interestingly, examination of the surrounding region using ChIP-seq revealed an adjacent H3K27me3 domain in the rabbit genome that is absent in other species and ends at the *TRAF6* promoter immediately adjacent to the site of CTCF enrichment (**Figure 6D**). These data imply that gain of H3K27me3 in the region of the *TRAF6* locus occurred in the common ancestor of rabbits and rodents, followed by loss of CTCF in the true rodent lineage that allowed extension of the H3K27me3 domain into the *TRAF6* promoter.

Our results suggest that shared features of the chromatin environment in germ and immune cells make them especially sensitive to *Traf6* regulatory mutations, and predict that other genes marked by newly emerging bivalent chromatin in germ cells may also have evolving immune functions. Indeed, we found that genes that recently gained species-specific germline bivalency, but not those with deeply conserved bivalency, are enriched for immune-biased inflammatory pathways such as NF-κB and TRIF/TLR4 signaling (**Figure 6E-F**) ^31^. Both newly-bivalent genes and genes in NF-κB/TLR4 pathways also had intermediate patterns of expression breadth, in contrast to housekeeping (mostly broad) or conserved bivalent (mostly tissue-specific) gene sets (**Figure 6G**). Notably, while overall expression levels of NF-κB and TRIF/TLR4 pathway genes were comparable between mouse and human in most tissues, their expression in testis was strongly depressed in mouse but not human (**Figure 6F**), similar to *Traf6.* Regulatory divergence at the *Traf6* promoter may thus be representative of coordinated expression evolution for multiple genes within the NF-κB/TLR4 inflammatory pathways, contributing to well-known differences in sensitivity to endotoxin challenge between humans and mice ^63,64^. Importantly, this set of genes appears to be dispensable for male fertility but impactful for immune function, providing a selective advantage and promoting evolutionary change.

## Discussion

Here, we uncovered a functional species difference in inflammatory gene expression by examining germline bivalency across multiple mammalian species. Replacing a short interval of noncoding promoter sequence in mice with orthologous human sequence was sufficient to fully humanize *Traf6* chromatin state in the testis without impacting spermatogenic development or fertility. Instead, the humanized promoter induced a strong immune phenotype, including humanization of the inflammatory response and increased cytokine production in macrophages. Together, our results point to immune and germ cell populations as sensitive detectors of regulatory sequence evolution at broadly-expressed genes with differing phenotypic requirements, such that tolerance of gene expression variation in germ cells balances selection for optimized immune function.

Our findings highlight an important fitness filter, where any regulatory change resulting in an advantageous somatic phenotype must also preserve gamete development. Thus, the pool of genes undergoing evolutionary shifts from broad to tissue-restricted expression is constrained by a requirement that their expression level in germ cells is not strongly linked to fertility. In this context, it is notable that spermatogenic cells are transcriptionally promiscuous, expressing most of the genome at some point in their development and therefore giving most new mutations an opportunity to affect germ cell function ^19–21,65^. However, many of these testis-expressed genes do not have a detectable impact on fertility when knocked out ^19–22,65^. Our results suggest that spermatogenic development may be optimized to tolerate variable levels of expression for most genes, allowing them to transmit mutations that confer an advantage to the organism as a whole. However, we note that our assessment of baseline fertility does not rule out potential germline vulnerabilities that might become apparent under conditions of physiological stress, aging, or environmental challenge.

In germ cells, evolution of *Traf6* regulation involved a tradeoff between CTCF binding and H3K27me3 deposition, where loss of CTCF binding permitted expansion of adjacent H3K27me3, generating a new bivalent domain and reducing expression. CTCF appears to act primarily as an activating transcription factor, rather than a looping factor or insulator, at the *Traf6* locus. In mESCs and germ cells, GC content and CpG island density are strong correlates of the bivalent state ^51–53^, but are antagonized by the presence of activating sequence ^25,66,67^. Notably, both inactive developmental genes in mESCs and inactive LPS-inducible genes in unstimulated macrophages have promoter chromatin features likely to make them transcriptionally sensitive to small changes in regulatory sequence: they are nucleosome-depleted, enriched for paused RNA polymerase II, and initiate low levels of transcripts even before stimulation ^68–71^. Altered binding of an activating transcription factor such as CTCF could tip the balance of expression in both cell types before affecting expression in other tissues, coordinating gain of bivalency at these genes in germ cells with their reduced expression in immune tissue. Bivalency in germ cells would become a readout of regulatory change that primarily impacts immune cells in the soma (**Figure 6H**).

Humans are known to be much more sensitive to endotoxin challenge than mice, requiring 500-fold less LPS per kilogram required to induce a similar level of cytokine response ^63,64,82^. Remarkably, a single small noncoding change was sufficient to recapitulate a portion of this phenotypic difference in our study. We also found evidence for reduced expression of other components of the NF-κB signaling pathway in the mouse lineage, suggesting that the species difference in endotoxin sensitivity is linked to cumulative regulatory changes at multiple genes in this pathway, with *Traf6* as a representative member of this set. In general, our analysis suggests an enrichment for immune function, and specifically for TLR and NF-κB signaling, among genes that have recently acquired germline bivalency.

Together, our results highlight the germ line as a critical filter in shaping regulatory evolution for broadly expressed genes, and indicate that divergence in germline bivalency can be used as a marker to identify genes with important and evolving roles in immunity. TRAF6 is also involved in other non-immune functions, including bone formation, tooth morphology, and development of hair follicles, sweat glands, and sebaceous glands ^46,72–78^, and has been implicated in tumorigenesis ^79–81^. Whether differential regulation of *Traf6* and other NF-κB pathway members impacts these functions, and the full effects of its differential role in mouse and human immune cells, remain open questions. By expanding our set of target loci and further comparing our humanized mice to additional human phenotypes, our model can uncover fundamental species differences in inflammatory responses and serve as an important system for studying human disease.

## Acknowledgments

We thank Tim Nottoli, Suxia Bai, and the Yale Genome Editing Core for generating the Traf6^h^ and Traf6^m^ mouse and ESC lines. We thank James Jusuf, Clarice Hong, and Anders Hansen for micro-C data visualization. We thank Julius Chapiro for rabbit samples and Mackenzie Noon for help with sgRNA design. We appreciate the work of the Yale Center for Genome Analysis in library preparation and high-throughput sequencing. This work was supported by the National Institute of Child Health and Human Development (R01HD098128 and R21HD110843), the National Cancer Institute (R21CA288677), an ACS-IRG pilot award from the Yale Cancer Center, and the Searle Scholars Program. Bluma Lesch, M.D., Ph.D. was supported by a Discovery Boost Grant, DBG-23-1150177-01-DMC, Grant DOI #10.53354/ACS.DBG-23-1150177-01-DMC.pc.gr.175431, from the American Cancer Society. B.J.L. is a Pew Scholar, supported by the Pew Charitable Trust.

## Author contributions

Conceptualization: KNG, KLM, GAR, BJL

Investigation: KNG, KLM, GAR, JA, DF, HY, EF, NRI, RL, KS

Formal analysis: KNG, KLM, GAR, JA, DF, HY, KS, BJL

Validation: KNG, KLM, GAR, JA, HY, KS

Visualization: KNG, KLM, GAR, BJL

Writing – original draft: KNG, KLM, BJL

Writing – review & editing: BJL

Resources: NJ, AW, BJL

Supervision: BJL Funding acquisition: BJL

## Methods

### Antibodies

The following antibodies were used in this study: anti-H3K4me3 (Abcam, ab8580); anti-H3K4me3 (Epicypher, 13-0041); anti-H3K27me3 (Abcam, ab6002); anti-H3K27me3 (Cell Signaling Technology, 9733); anti-H3 (Abcam, ab1791); anti-CTCF (Active Motif, 61932); anti-GFP (Abcam, ab13970); anti-TRAF6 (Abcam, ab33915); 488-Donkey anti-Chicken (Jackson ImmunoResearch Labs, 703-545-155).

### Mice

These studies were approved by the Yale University Institutional Animal Care and Use Committee under protocol 2023-20169. All animals used in these studies were maintained and euthanized according to the principles and procedures described in the National Institutes of Health Guide for the Care and Use of Laboratory Animals. Mice were kept under standard conditions with access to food and water *ad libitum* and all experiments were conducted in compliance with the Animal Welfare Act. For fertility testing, homozygous Traf6^h/h^ or wild type control males were co-housed with wild type females for six months starting at three months of age and litter sizes and dates were recorded at weaning.

### mESC culture

Mouse embryonic stem cells were grown on 0.1.% gelatin (Millipore ESE-006-B) and mouse embryonic fibroblast (MEF) feeder cells (Gibco MitC-treated CF-1 MEF A34959) at 37C and 5% CO2 in a humidified incubator in embryonic stem cell media [DMEM (Gibco 11965-092) with 15% fetal bovine serum (Sigma-Aldrich F2442), Penicillin-Streptomycin (1:100; Gibco 15140-122), GlutaMax (1:100; Gibco 35050-061), MEM NEAA (1:100; Gibco 11140-050), Sodium Pyruvate (1:100; Gibco 11360-070), HEPES (1:40; Gibco 15630-080), β-mercaptoethanol (1:125,000; Sigma-Aldrich M6250), and LIF (1:10,000; Millipore ESG1106)]. Media was changed daily and cells were passaged every 2-3 days with StemPro Accutase (Gibco A11105-01). Growing cells were tested for mycoplasma every 1-2 months to ensure no contamination.

### Generation of transgenic mice and mESCs

Transgenic mouse embryonic stem cells and mice were generated at the Yale Genome Editing Center. Traf6^h^-GFP and Traf6^m^-GFP heterozygous mESC lines were generated by CRISPR-mediated homologous recombination on a C57BL/6 genetic background. The transgenes and strategy are illustrated in **Figure S1**: for Traf6^h^-GFP, 1475 base pairs of human sequence (chr11: 36,509,376-36,510,850 (hg38) replaced 1171 base pairs of orthologous mouse *Traf6* noncoding region (chr2:101,508,481-101,509,651 (mm39)), and an *Egfp* coding sequence was inserted downstream. For Traf6^m^-GFP, only the *Egfp* sequence was inserted in the same location. Positive clones were genotyped by PCR at 5’ and 3’ ends and karyotyped to confirm a normal chromosome complement. The neomycin selection cassette was removed by transient transfection of a Flp recombinase followed by selection and genotyping for neomycin-negative clones.

Traf6^h^-GFP and Traf6^m^-GFP mice were generated by blastocyst injection of Traf6^h^-GFP mESCs to generate chimeras, and chimeras with germline transmission were used to generate founders. The neomycin cassette was removed by crossing in an Actb-Flp transgene. After confirmation of neomycin excision, the Actb-Flp transgene was crossed out to generate neo-transgenic mice. Traf6^h^-GFP and Traf6^m^-GFP mice were backcrossed to C57BL/6 (Jackson Labs 000664) for at least six generations. Traf6^h/h^ mice were generated by the Yale Genome Editing Center by zygotic CRISPR injection to delete the *Egfp* coding sequence from the neo-Traf6^h^-GFP mouse line. The excision leaves a 70 base pair scar in the first intron of *Traf6*. Founders confirmed to carry the *Egfp* deletion were used to establish the Traf6^h^ line, backcrossed to C57BL/6 for at least four generations, and crossed together to generate Traf6^h/h^ homozygous mice.

### CRISPR editing in mESCs

#### Single-guide RNA (sgRNA) plasmid creation

sgRNAs were designed to target the NGG sites nearest to and flanking the identified CTCF binding domain within the swapped human promoter sequence (**Table S6**). sgRNAs were cloned into lentiCRISPR v2 plasmid with a Cas9 and puromycin selection cassette (Addgene, 52961; RRID: Addgene_52961) as previously described ^83,84^. Briefly, plasmid was digested with BsmBI-v2 (NEB, R0580) at 55°C for 1 hour. Digestion was separated on 1% agarose gel and band was extracted using QIAquick Gel Extraction Kit (28704, Qiagen) according to manufacturer instructions. sgRNA oligos were phosphorylated and annealed using T4 PNK (NEB, M0201S) and T4 Ligation Buffer (NEB, B0202S). Oligos were diluted 1:100 and ligated with digested lentiCRISPR v2 plasmid using T4 Ligase (NEB, M0202S) for 1 hour at room temperature. Plasmid was transformed into NEB stable competent *E. coli* (C3040H, NEB) according to manufacturer instructions. Bacteria were plated on LB plates with ampicillin (1:100) and grown overnight at 37C. Individual colonies were cultured in LB-ampicillin media shaking overnight at 37C. Plasmids were isolated using Qiagen QIAprep Spin Miniprep Kit (27106, Qiagen) according to manufacturer instructions.

#### Transfection of Traf6^h^-GFP mESCs

One day prior to transfection, Traf6^h^-GFP embryonic stem cells were plated at a low density (2.5 x 10^5^ cells) on 0.1% gelatin (Millipore, ES-006-B). Cells were transfected with sg1 alone or both sgRNA pairs using the Lipofectamine 3000 Transfection Kit (Invitrogen, L3000015) according to manufacturer instruction. After 14-18 hours, cells were passaged into 10 cm polystyrene culture dishes coated with 0.1% gelatin and MEF feeder cells (Gibco, A34959). Beginning 24 hours after transfection, cells were selected in media with 1.5 ug/mL puromycin for 48 hours, including a non-transfected control for each cell line. After selection, transfected cells were cultured for 3-6 days to allow growth to confluence. Sanger sequencing was performed to confirm editing.

### Generation of single cell suspensions

#### Whole testis

Testes from adult male mice were removed from the scrotum, decapsulated from the tunica albuginea, and incubated in 0.75mg/mL collagenase (Gibco, 17104-019; Thermo Fisher Scientific, Inc.) in Dulbecco’s Modified Eagle Medium (DMEM; Gibco 11965-09L; Thermo Fisher Scientific, Inc.) at 37°C for 10 minutes. DMEM was added to dilute collagenase and samples were centrifuged for 5 minutes at 400 x g at room temperature. The supernatant was discarded and samples were washed in DMEM. Samples were resuspended in 0.05% trypsin-EDTA (Gibco 25300-054; Thermo Fisher Scientific Inc.) with DNAse (1:10,000; Stem Cell Technologies, 07900). The reaction was quenched with 10% Cosmic Calf Serum (Sigma-Aldrich, C8056) in DMEM (CCS-DMEM) and centrifuged. Samples were resuspended in CCS-DMEM and filtered through a 100-micron filter. Cell concentration was determined by hemocytometer.

#### Bone Marrow

Tibia and femurs were harvested and placed in ice cold media containing 2% FBS. Connective tissue and muscle were removed and bones were cut at each end to expose bone marrow. Bones were flushed with ice cold PBS + 2% FBS with a 25G needle. The suspension was disrupted with a 1ml pipette and filtered over a 70 um filter. The flow-through was collected and centrifuged at 300–400 × g for 5 minutes at 4°C. Red cell lysis was performed as needed.

#### Spleen

Spleens were harvested and placed in cold PBS supplemented with 2% FBS. Using a syringe plunger, spleens were gently mashed through a 70 μm cell strainer to create a single-cell suspension. The flow-through was collected and centrifuged at 300–400 × g for 5 minutes at 4°C. Following mechanical disruption, red cell lysis was performed for 5 minutes at room temperature.

#### Thymus

Thymuses were harvested and placed in cold PBS supplemented with 2% FBS. Using a syringe plunger, tissues were gently mashed through a 70 μm cell strainer to create a single-cell suspension. The flow-through was collected and centrifuged at 300–400 × g for 5 minutes at 4°C.

#### Blood

30-50 ul of blood was collected by retro-orbital bleed. Red cell lysis was performed for 5 minutes at room temperature. Lysis step was quenched with 4x volumes of cold PBS supplemented with 2% FBS. This step was repeated until pellets were no longer red.

#### Liver

Livers were excised, weighed, and manually chopped using a razor blade. For immune cell isolation, chopped tissue was digested in digestion buffer containing 1% Bovine Serum Albumin, 2mg/mL collagenase IV, and 0.1mg/mL DNAse I for 30 minutes at 37C, 200rpm. Post-digestion, livers were mashed using a 70 um filter, washed with 1X PBS, and run on a 33%/66% Percoll gradient, and interphase cells were collected.

### Histological staining

Whole testes were fixed in Hartman’s Fixative (Sigma-Aldrich, H0290) overnight at 4°C, then transferred to 70% Ethanol. Testes were cross-sectioned followed by dehydration and paraffin embedding. Paraffin sections were mounted on slides and stained with hematoxylin and eosin to visualize DNA and cell structure. Testis sections were imaged at 20x or 40x magnification using a Zeiss AxioImager M1 with attached camera Zeiss AxioCam MRc in brightfield.

### Immunofluorescence

For immunofluorescence staining, tissue sections were fixed for 8 minutes in 4% paraformaldehyde (PFA; Sigma-Aldrich 158127) in PBS containing 0.2% Triton X-100 (PBS-T; AmericanBIO, AB02025). Sections were washed in PBS-T, then incubated in blocking buffer (3% BSA (Sigma-Aldrich, A9647), 5% NDS (Jackson ImmunoResearch Labs, 017-000-121) in PBS-T) for 15 minutes. Then, primary antibody (chicken anti-GFP, Abcam ab13970) diluted 1:1000 in blocking buffer was added to sections for 15 minutes at room temperature (RT). After washing sections three times in PBS-T, sections were incubated in secondary antibodies (488-Donkey anti-chicken, Jackson ImmunoResearch Labs, 703-545-155) diluted 1:200 in blocking buffer for 10 minutes at RT. Sections were washed in PBS-T, then mounted in 90% glycerol (JT Baker, 2136-01) in PBS plus 2.5 mg/ml *p-*Phenylenediamine (Acros Organics, 417481000). Tissue sections were imaged on a Zeiss AxioImager Z1 microscope with Apotome.2 attachment, Plan-APOCHROMAT 20X/0.8 air objective, Zeiss Axiocam 506 mono camera, and Zen software (v3.0; Zeiss).

### Flow cytometry

#### Round spermatid isolation

Round spermatids were isolated by flow cytometry as previously described ^85^. Briefly, dissociated testis cells were incubated at 37°C with 2 µl/ml Vybrant DyeCycle Green (Life Technologies, V35004) for 20 minutes. Round spermatids were sorted based on 1C DNA content and size using a 2-laser sorter (Bio-Rad S3e with 488nm and 561nm lasers) gated on single cells.

#### GFP quantification

Cells were harvested using StemPro Accutase (Gibco, A11105-01), resuspended in DMEM and kept on ice. GFP expression was quantified using a BioRad S3e Cell Sorter (BioRad). Laser settings were established for each replicate using the Traf6^m^-GFP cell line and kept constant for all acquisitions. Mean GFP fluorescence of each cell line was calculated for each replicate (n = 3-5). Tukey’s multiple comparison test was performed to evaluate each pairwise relationship.

#### Immune profiling

Peripheral blood, spleen, thymus, liver, and bone marrow were harvested according to standard protocols as described above. Dissociated cells were stained with extracellular antibodies in FACS buffer (0.5% FBS, supplemented with EDTA, and 1× PBS without Mg2+/Ca2+) at 4°C for 30 min followed by fixation with Cytofix/Cytoperm Fixation/Permeabilization from BD Biosciences. Staining reagents included BV421-CD3e (17A2, Biolegend), BV510-CD62L (Mel-14, Biolegend), BV605-CD11b (M1/70, Biolegend), BV711-Ly6c (HK1.4, Biolegend), BV-785-CD44 (IM7, Biolegend), FITC-NK1.1 (PK136, Biolegend), PerCPCy5.5-CD4 (GK1.5, Biolegend), PE-CD25 (PC61, Biolegend), PE-Dazzle 594-CD45 (30-F11, Biolegend), PE-Cy7-CD11c (N418, Biolegend), APC-CD19 (6D5, Biolegend), BUV395-CD8a (53-6.7, BD), BUV737-Ly6g (1A8, BD), Live/Dead fixable Near IR (Invitrogen).

For peripheral blood, spleen, and bone marrow, cell populations were defined as a fraction of live cells (CD45+) using the following marker combinations: total myeloid (CD11b+); neutrophil or granulocyte (CD11b+ Ly6C+ Ly6G+); monocyte or macrophage (CD11b+ Ly6C+ Ly6G−); total T cells (CD3+); CD4+ T cells (CD3+ CD4+); CD8+ T cells (CD3+ CD8+); B cells (CD19+); NK cells (NK1.1+). For thymus, total myeloid (CD11b+) and B cells (CD19+) were defined in the same way as a fraction of live cells (CD45+), and remaining cell populations were defined as a fraction of non-myeloid, non-B (CD11b− CD19−) cells using the following marker combinations: CD4+ single positive (CD4+); CD8+ single positive (CD8+); double negative (CD4− CD8−); double positive (CD4+ CD8+).

### LPS challenge and ELISAs

LPS from *Escherichia coli 055:B5* was prepared at a stock concentration of 10mg/mL in PBS, aliquoted, and stored at −80C. Upon challenge, aliquots were further diluted in sterile PBS, and mice were treated via intraperitoneal (i.p.) injection with an LD_50_ dose calculated based on body weight and sex (6mg/kg for females, 7.5mg/kg for males) ^86^. Mice were grouped-housed, given *ad libitum* access to food and water throughout challenge, and acclimated to the experimental space for a minimum of 2 days prior to challenge. For plasma collection, mice were bled via retro-orbital route under isofluorane at 0 hours, 2 hours, and 6 hours post-treatment, and plasma was immediately isolated using lithium heparin centrifugation tubes, and stored at −80C. For quantification of plasma cytokines by ELISA, anti-mouse IL-6 (eBioscience 14-7061-85) and anti-mouse/rat TNFα (Invitrogen 14-7423-85) were used as captures antibodies. Biotinylated anti-mouse/rat IL-6 (BD Pharmingen 554402) and biotinylated anti-mouse/rat TNFα (eBioscience 13-7341-85) were used as detection antibodies. For both ELISAs, Streptavidin HRP (BD Pharmingen 554066) was used as secondary detection antibody, and plates were developed using TMB substrate reagent (BD 555214) and stopped with 1N hydrochloric acid (VWR E447-1L). Statistical comparisons were performed with multiple unpaired t-tests for each tissue individually. Three independent biological replicates (n=5, 3, and 2 for Traf6^h/h^ and n=5, 3, and 1 for control) were performed. Controls were either wild type littermates of Traf6^h/h^ experimental animals or C57BL/6J animals obtained at 5 weeks of age from Jackson Laboratories (strain #000664) and housed for a minimum of 4 weeks in the same colony as experimental animals to acclimatize the microbiome.

### Bone marrow derived macrophages (BMDM)

Bone-marrow cells were flushed from the femurs of Traf6^h/h^ or control mice using a syringe and supplemented RPMI media (RPMI 1640 + 10% FBS + 1% Penicillin/Streptomycin). Marrow cells were strained through a 70 micron filter and centrifuged at 1800rpm for 5 minutes at 4C. Red blood cell lysis was performed using ACK Lysis Buffer (Gibco) for 5 minutes at room temperature, then samples were washed with supplemented RPMI media. After counting, 5-10×10^6^ cells per femur were plated in 10cm bacterial petri dishes, in 15mL of RPMI-LCCM (supplemented RPMI containing 30% L929 cell cultured media), and cells were placed at 37C 5% CO2 for differentiation. On the third day of culture, each dish was supplemented with an additional 10mL of RPMI-LCCM. On the sixth day of culture, each plate was washed a minimum of 2 times with sterile PBS. Differentiated BMDMs were then detached using Cell Stripper (Gibco), plated at a density of 2×10^5^ cells per well in a 24 well-plate, and allowed to adhere overnight in 500ul of supplemented RPMI media. The following day, stimulations were performed in 300ul of supplemented RPMI media per well, containing either 1ng/mL LPS from *Escherichia coli* 055:B5, or recombinant mouse TNFalpha (Biolegend 575208). After stimulation, supernatants were collected and centrifuged briefly before storing at −80C. ELISAs were performed using the same protocol as described above for mouse plasma samples.

### RNA isolation and reverse transcription

Dissected tissues were placed in 1ml Trizol and disrupted by drawing up and down in an 18G followed by a 21G needle and syringe. Aliquots of mESCs or bone marrow cell suspensions collected as described above were centrifuged at 6000 xg for 4 minutes at 4C, resuspended in 500 uL Trizol (Ambion, 15596018, Thermo Fisher Scientific, Inc.) and disrupted by drawing up and down through at 21G needle and syringe. 50 uL 1-bromo-3-chloropropane (BCP; Sigma-Aldrich, B9673) per 500 uL of Trizol was added and samples were centrifuged at 12000 xg for 15 minutes at 4°C. The aqueous layer was applied to gDNA eliminator columns supplied in the Qiagen RNeasy Plus Mini kit (#74134) and RNA from the resulting flow-through was extracted using the RNeasy Plus Mini kit according to the manufacturer’s instructions. Samples were processed in batches of 1-5 in order of collection and stored at −80°C. Total RNA was used for RNA-seq library preparation (see below) or reverse transcription. Reverse transcription was performed with oligo dT primers and SuperScript III reverse transcriptase (Invitrogen, 18080-051, Thermo Fisher Scientific, Inc.) according to the manufacturer’s instructions. Reaction mixtures (20 uL) were incubated in a thermocycler for 50 minutes at 50°C followed by 5 minutes at 85°C to stop the reaction. cDNA was diluted in nuclease-free water as necessary prior to qPCR.

### qPCR

Real time quantitative PCR was performed in 20 uL reaction volume (1-4 uL diluted cDNA, 10 uL Power SYBR Green PCR Master Mix (Applied Biosystems, 4367659, Thermo Fisher Scientific, Inc.), 5.6-8.6 uL nuclease-free water, and 0.4 uL of 10 uM forward and reverse primer mixture). Reactions for each target were performed in triplicate in a 96 well plate using a QuantStudio 3 Real-Time PCR System (Applied Biosystems, Thermo Fisher Scientific, Inc.) with standard cycling conditions (Hold stage (x1): 50°C for 2 minutes, 95°C for 10 minutes; PCR stage (x40): 95°C for 15 seconds, 60 °C for 1 minute, Melt curve stage: 95°C for 15 seconds, 60°C for 1 minute, 95°C for 15 seconds). For RT-qPCR, relative expression was calculated using the comparative Ct method with *Actb* as an internal control. Primer sequences are provided in **Table S6**.

For ChIP-qPCR, enrichment at a specific locus was calculated using comparative Ct, normalized to the negative control locus (mm10 chr19:51758385-58758494) and to the endogenous mouse allele when indicated. Statistical comparisons were performed by Welch’s t-test or 1-sample t-test relative to an expected value of 1 as appropriate. Primer sequences are provided in **Table S6**.

### Chromatin immunoprecipitation (ChIP)

For histone modification ChIP, ChIP was carried out as previously described ^31,87^. Dissociated testis or mESCs (0.2-5 x 10^6^ cells) were cross-linked with 1% methanol-free formaldehyde at room temperature for 10 minutes, then quenched with 2.5M glycine at room temperature for 10 minutes. Fixed cells were centrifuged at 6,000 xg for 3 minutes at 4°C. Cells were washed with cold PBS twice, resuspended in 100 μL ChIP lysis buffer (1% SDS, 10mM EDTA, 50mM Tri-HCl at pH 8.1) and snap frozen in liquid nitrogen. Dynabeads (Invitrogen, 00821318) were prepared by mixing 10 μL of beads with 100 μL block solution (0.5% BSA in PBS). Beads were washed twice with 150 μL of block solution and resuspended in 30ul block solution. 0.5 μL of H3K4me3 (Abcam, ab8580) or 1 μL of H3K27me3 (Abcam, ab6002) was added to each aliquot of beads and incubated for 8 hours rotating at 4°C. Fixed frozen cells were thawed on ice and diluted to a total volume of 300 μL per sample with ChIP dilution buffer (0.01% SDS, 1.1% Triton X-100, 1.2mM EDTA, 167mM NaCl, 16.7mM Tris-HCl at pH 8.1). Cells were split into 150 μL aliquots in 0.5 ml Eppendorf tubes and sonicated at 4°C for 30 cycles (30 seconds on/off) using a Bioruptor (Diagenode). Aliquots of the same samples were pooled and centrifuged at 12,000 x g for 5 minutes. Supernatant was collected in a new 1.5 mL Eppendorf tube and combined with 600 μL of dilution buffer and 100 μL protease inhibitor cocktail (Roche, 11836153001). 50 μL of each sample was set aside as input. To all other samples, appropriate antibody-bound Dynabeads were added and mixtures were incubated overnight at 4°C, rotating. Beads were washed twice with low-salt immune complex wash buffer (0.1% SDS, 1% Triton X-100, 2mM EDTA, 150mM NaCl, and 20mM Tris-HCL at pH 8.1), twice with LiCl wash buffer (0.25M LiCl, 1% NP40, 1% deoxycholate, 1mM EDTA, and 10mM Tris-HCl at pH 8.1), and twice with TE (1 mM EDTA and 10mM Tris-HCl at pH 8.0). Bound DNA was eluted twice with 125 μL elution buffer (0.2% SDS, 0.1M NaHCO3, and 5mM DTT in TE) at 65°C. Eluted bound DNA and input samples were incubated for 8 hours at 65°C to reverse crosslinking followed by 2 hours at 37°C with 0.2 mg/mL RNAse A (Millipore, 70856-3), and 2 hours at 55°C with 0.1 mg/mL Proteinase K (NEB, P8107S).

For CTCF, ChIP was carried out as previously described ^88^ with some modifications. For each ChIP reaction, 3 x 10^6^ cells were cross-linked in 1% methanol-free formaldehyde at room temperature for 10 minutes, then quenched with 0.114 M Glycine. Fixed cells were resuspended in 300 uL High Salt Lysis/Sonication Buffer (800 mM NaCl, 25 mM Tris pH 7.5, 5 mM EDTA, 1% Triton X-100, 0.1% SDS, 0.5% sodium deoxycholate, 1 x protease inhibitor cocktail) and frozen in −80°C. 20 uL Dynabeads Protein G (Thermo Fisher Scientific) was first incubated with 1 μL of CTCF antibody (Active Motif, 61932) in 20 uL Chromatin Dilution Buffer (25 mM Tris pH 7.5, 5 mM EDTA, 1% Triton X-100, 0.1% SDS, 1 x protease inhibitor cocktail) at room temperature for 3 hours with rotation. Fixed cells were transferred to a Bioruptor Pico Microtube (Diagenode) and sonicated for 10 cycles (30 seconds on/off) in Bioruptor (Diagenode) at 4°C. 1 mL Chromatin Dilution Buffer was added to sonicated cells, and samples were spun at 13600 x g for 30 minutes at 4°C to pellet insoluble material. 65 uL soluble chromatin was set aside as input. Dynabead/antibody mix was added to the soluble chromatin and incubated at 4°C overnight with rotation. Then, beads were washed with 1 mL Wash Buffer A (140 mM NaCl, 50 mM HEPES pH 7.9, 1 mM EDTA, 1% Triton X-100, 0.1% SDS, 0.1% sodium deoxycholate), Wash Buffer B (500 mM NaCl, 50 mM HEPES pH 7.9, 1 mM EDTA, 1% Triton X-100, 0.1% SDS, 0.1% sodium deoxycholate), and Wash Buffer C (20 mM Tris pH 7.5, 1 mM EDTA, 250 mM LiCl, 0.5% NP-40 Alternative, 0.5% sodium deoxycholate) once each, and TE Buffer (10 mM Tris pH 7.5, 1 mM EDTA) twice. Bound DNA was eluted twice with 100 uL Elution Buffer (10 mM Tris pH 7.5, 1 mM EDTA, 1% SDS) at 65°C for 5 min. Eluted DNA and input samples were incubated with 4 ug RNase A (Millipore Sigma) at 65°C overnight to reverse cross-linking and digest any contaminating RNA. ChIP and input samples were further digested with 40 ug Proteinase K (NEB) at 45°C for 2 hours.

DNA from all ChIP and input samples was concentrated using Zymo ChIP DNA Clean & Concentrator kit (Zymo Research, D5201) according to manufacturer instructions.

### Re-ChIP

Whole testes were dissociated to a single cell suspension as described above and 6 x 10^6^ mixed testicular cells were used as input. Re-ChIP was performed using the Re-ChIP-IT kit (Active Motif, 53016) according to the manufacturer’s instructions, using 2 ul anti-H3K4me3 (Abcam, ab8580) for the first IP and 2 ul anti-H3K27me3 (Abcam, ab6002) for the second IP.

### Library preparation and sequencing

Purified ChIP and input DNA were used for sequencing library preparation. RNA-seq libraries were prepared from total RNA using poly-A selection with the KAPA mRNA HyperPrep Kit (Roche, 08098123702). ChIP-seq libraries were prepared using either the KAPA HyperPrep Kit (Roche, KK8504) or the Watchmaker DNA Library Prep Kit (7K0103-096). Samples were sequenced on an Illumina HiSeq6000 machine with paired 150 base pair reads to a depth of approximately 30 million reads.

### ChIP-seq data analysis

#### Rat and rabbit ChIP-seq data processing

Data were filtered for high-quality reads using the fastq_quality_filter tool from FASTX-Toolkit with parameters -p 80 -q 20. Filtered libraries were aligned to the rat (rn6) or rabbit (oryCun2) genome assemblies using Bowtie2 ^89^ (--end-to-end --sensitive). Aligned reads were filtered to get uniquely mapping reads without PCR duplicates using SAMtools ^90^ (-q 1 -f 3 -F 256). Peaks were called using MACS2 ^91^ with default parameters. Peak intersections were evaluated using BEDTools ^92^. Bigwig files were generated using the bamCoverage tool from deepTools and used for visualization in the UCSC genome browser.

#### Genome-wide ChIP-seq signal quantitation

For quantitation of ChIP-seq signal shown in **Figure 1A**, round spermatid ChIP-seq libraries were obtained from published datasets (human, rhesus, mouse, opossum, bull: ^31,93^ GEO: GSE68507 and GSE49621). Data were filtered for high-quality reads using the fastq_quality_filter tool from FASTX-Toolkit with parameters -p 80 -q 20. Overrepresented adaptor sequences were trimmed using CutAdapt and reads were aligned using Bowtie2 ^89^ (--end-to-end --sensitive). PCR duplicates were removed using Picard MarkDuplicates ^94^ (-REMOVE_DUPLICATES true). Aligned .bam files were converted to bedgraph files using deepTools bamCoverage ^95^ (--normalizeUsing CPM --outFileFormat bedgraph), with the effective genome size parameter calculated using library read length and unique-kmers.py in khmer ^96–98^. To obtain signal strength information in promoters, promoter intervals were generated using bioMart (Ensembl v.113). For opossum, a gene .gtf file was obtained from UCSC bigZips (monDom5.ensGene.gtf). A single transcriptional start site extracted from the longest transcript was chosen for each unique gene ID and a 2kb window centered on the TSS was defined. Read signal from bedgraph files was measured within promoter intervals using bedTools map ^92^ (-c 4 -o sum -null 0 -F 0.5). The summed signal within promoter intervals was averaged across biological replicates, except for bull, where only 1 biological replicate was available. To more clearly visualize the data, a Yeo-Johnson power transformation was applied. First, H3K4me3 and H3K27me3 signal outliers were removed manually by visualizing the data and establishing appropriate signal thresholds. On average, this step resulted in retention of 0.995 of the original regions. Next, the transformation was performed using scipy.stats.yeojohnson ^99,100^. Transformed signal was plotted using Seaborn jointplot ^101,102^.

#### Metagene plots

To determine histone modification signal at CTCF peaks, bigwig files were generated from the .bdg files used above using Kent Tools bedGraphToBigWig (--version 2.8) and replicates were averaged using deepTools bigwigAverage (--version 3.5.5). CTCF narrowPeak files were downloaded from ENCODE (mouse accession: ENCFF443TPY; human accession: ENCFF644JKD, ENCFF767LMP) ^35,36^. For human data, the two peak files were intersected (bedtools intersect) to define biologically replicated CTCF peaks (v2.30.0). Mean histone modification ChIP-signal was calculated at CTCF peaks using deepTools computeMatrix reference-point (-a 2000 -b 2000 --referencePoint center –skipZeros). This resulting matrix was then plotted using plotProfile.

### RNA-seq data analysis

RNA-seq libraries were aligned using HISAT 2.2.1 ^103^ with Ensembl ^104^ release 104 as a reference and data output type. Raw count values and FPKM were obtained using StringTie 2.1.4 ^105^ with the -e and -G options and Ensembl GRCM39 release 104 reference files. Differentially expressed genes between Traf6^h^-GFP and Traf6^m^-GFP samples were called using DESeq2 ^106^ with n=3 biological replicates.

### Pan-tissue gene expression

Tau metrics were calculated from polyA+ RNA-seq FPKM values. Expression data was obtained from ENCODE with identifiers listed in **Table S1**. For mouse, all tissues were sampled from 2-month old male B6NCrl mice. For human, all tissues were sampled from male adults. Tau values were calculated using tspex ^33^ and expression values generated by ENCODE ^36^. To visualize expression across tissues, continuous FPKM values were binned into categorical expression levels. For each tissue, an FPKM cutoff was determined using the 95^th^ quantile. All genes with expression over this cutoff were set to the 95^th^ quantile value in order to expand the dynamic range of the middle bins. Next, for each tissue, the filtered FPKM values were assigned an expression bin using the “cut” function in pandas, n=10, with 0 equaling lowest expression and 9 equaling highest expression in the tissue. The FPKM table was then subset to genes of interest, and the categorical expression level was plotted as a heatmap using the function “clustermap” in Seaborn. The “housekeeping” gene set was defined as the 100 genes with lowest tau score. Core poised genes and species-specific bivalent genes were defined from supplementary tables from ^31^. To facilitate visualization given the large size of the species-specific bivalency lists (human=796, mouse=490), a random sample of 300 species-specific bivalent genes was plotted for both mouse and human. NF-κB signaling and TRIF signaling genes were defined from GO categories GO_0043123 and GO_0035666, respectively. Expression level of a given gene is shown across 11 human tissues and 12 mouse tissues.

### Micro-C analysis

3D genome structure maps (**Figure S6**) of the *Traf6* region in human H1-hESC cells and mouse ES cells were plotted using Micro-C data. The human dataset was obtained from ^107^ (4DN accession number: 4DNES21D8SP8), and the mouse dataset was a merge of several existing mESC Micro-C datasets compiled at GSE286495 and reported in ^108^.

## Data availability

RNA-seq and ChIP-seq data is available at the NCBI GEO database under accession GSE297884.

## Supplemental Figure Legends

**Figure S1. Species differences in *Traf6* bivalency in sperm and generation of *Traf6* knock-in alleles for this study.**

**A,** Genome browser tracks showing H3K4me3 and H3K27me3 ChIP-seq signal in mouse epididymal sperm or human ejaculated sperm. Data from GSE42629 (mouse) or GSE15690 (human).

**B,** Schematic of generation of the Traf6^h^-GFP and Traf6^h^ alleles. Traf6^h^ was generated by removing the GFP coding sequence from Traf6^h^-GFP. A short piece of the En2 splice acceptor is retained, and 70 base pairs are deleted from the first *Traf6* intron.

**C,** Representative genotyping for litters carrying Traf6^h^-GFP or Traf6^h^ alleles.

**D,** Schematic of generation of the Traf6^m^-GFP allele.

**E,** Representative genotyping for litter carrying Traf6^m^-GFP.

**Figure S2. Specificity of human and mouse *Traf6* primers and bivalency at the Traf6^m^-GFP allele.** ChIP-qPCR for H3K4me3 and H3K27me3 using primers specific to the endogenous mouse or substituted human *Traf6* promoter sequence.

**A,** Dissociated testis tissue from Traf6^h^-GFP mice.

**B,** Sorted germ cells (round spermatids) from Traf6^h^-GFP mice.

**C,** Dissociated testis tissue from Traf6^m^-GFP mice. No human sequence is present and there is no amplicon from the human primers.

Data in **A** and **B** are also shown in **Figure 1F** and **1G** as a ratio between human and mouse alleles. Y-axis represents fold change relative to a negative control locus (mm10 chr19:51758385-58758494). N=3 biological replicates; plots show mean +/− s.e.m.

**Figure S3. Flow cytometry gating strategy for immune populations in Figures 4 and S4.**

**A,** Gating strategy for non-bone marrow cell populations. Nested gates are indicated by colored boxes.

**B,** Gating strategy for bone marrow cell populations. Nested gates are indicated by colored boxes.

**Figure S4. Additional immune profiling of Traf6^h/h^ animals.**

**A** to **D,** Relative fractions of circulating immune populations in spleen **(A)**, lung **(B)**, bone marrow **(C)**, and liver **(D)** assessed by flow cytometry. APC, antigen presenting cell. *p<0.05, **p<0.01, ***p<0.001, ****p<0.0001, unpaired t-test.

**E** and **F,** Mean GFP fluorescence intensity for different populations of immune cells in bone marrow (**E**) and in the intestinal epithelial lymphocyte compartment (**F**) from Traf6^h^-GFP or Traf6^m^-GFP mice. *p<0.05, **p<0.01, ***p<0.001, ****p<0.0001, unpaired Welch’s t-test.

**G,** Immunofluorescence for GFP (magenta), pan-immune marker CD34 (cyan), and DAPI (white) in small intestine (jejunum) epithelium of Traf6^m^-GFP and Traf6^h^-GFP mice. The same images are shown in Figure 4J; here, DAPI stain is added to show nuclei. Scale bar, 20um.

**H,** Mean GFP signal in Traf6^h^-GFP epithelial cells, normalized to average Traf6^m^-GFP signal across n=3 biological replicates. *p<0.05, unpaired t-test.

**I,** Relative expression (RT-qPCR) of *Traf6* transcript assayed by RT-qPCR in different bulk tissues of Traf6^h/h^ compared to control. N=3-10, *p<0.05, ****p<0.0001, one-sample t-test with expected value of 1.

**Figure S4. Contact maps for the *Traf6* locus in human and mouse ESCs.** Contact frequencies from Micro-C data in human (**A**) or mouse (**B**) ESCs. Box shows syntenic regions and dashed line indicates location of the *Traf6* promoter along the x-axis.

**Figure S5. Conservation and divergence of regulation at *Traf6* and immune genes.**

**A** and **B,** Mean H3K4me3 and H3K27me3 ChIP-seq signal from post-meiotic male germ cells (round spermatids) in ENCODE testis CTCF peaks in human (**A**) or mouse (**B**).

**C,** Diagram of the human *TRAF6* and mouse *Traf6* loci showing the swapped orthologous region, site of CTCF enrichment according to ENCODE, and putative divergent CTCF binding site. There is no predicted CTCF binding site under the ENCODE peak region in either species. Locations of PCR primers for Figure 5E and S6D are shown.

**D,** CTCF ChIP-qPCR at the ENCODE-annotated CTCF peak in mouse and humanized Traf6 promoter sequences in Traf6^h^-GFP mESCs. *p<0.05, **p<0.01, unpaired t-test, n=3 biological replicates.

**E,** Heatmap of gene expression decile across mouse tissues for housekeeping (n=100), conserved bivalent (n=351), human-specific bivalent (n=300, random sample of 490) genes, and for genes in the GO classes “Positive regulation of canonical NF-κB signal transduction” (GO:0043123, n=326) and “TRIF-dependent Toll-like receptor signaling pathway” (GO:0035666, n=13).

## Supplemental Tables

**Table S1.** Public datasets used to obtain gene expression values in mouse and human tissues.

**Table S2.** Public datasets used to calculate histone modification enrichment values in Figure 1C.

**Table S3.** Differential expression analysis of RNA-seq data for Traf6^h^-GFP vs. Traf6^m^-GFP in Figure 2D.

**Table S4**. Peak calling for ChIP-seq data for H3K4me3, H3K27me3, and CTCF in rat.

**Table S5.** Peak calling for ChIP-seq data for H3K4me3, H3K27me3, and CTCF in rabbit.

**Table S6**. Primers used in this study.

