## Supplementary figures and images for "Complementary constraints in germ and immune cells shape evolution of gene regulation and phenotype"

### Supplemental Figures

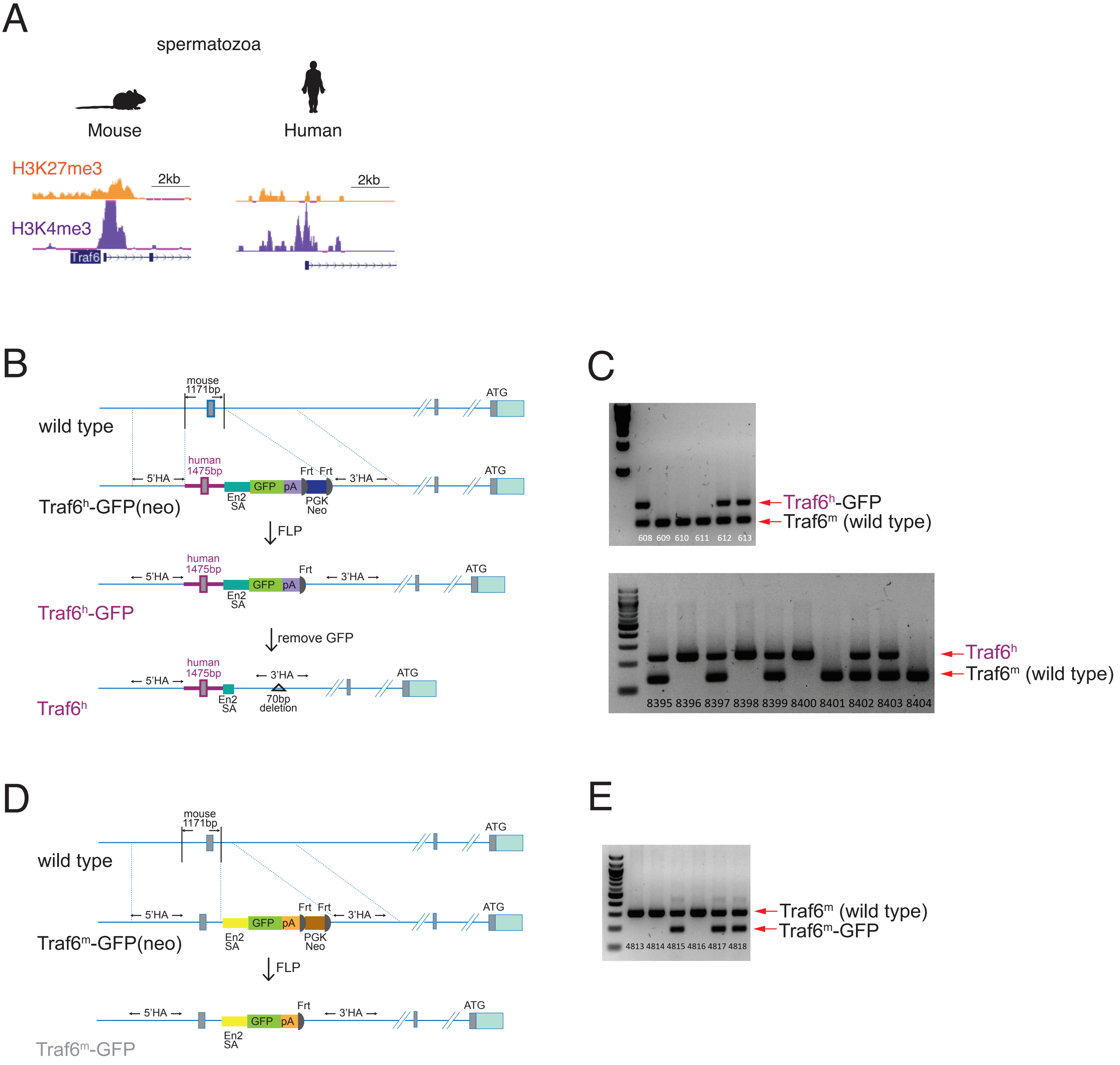

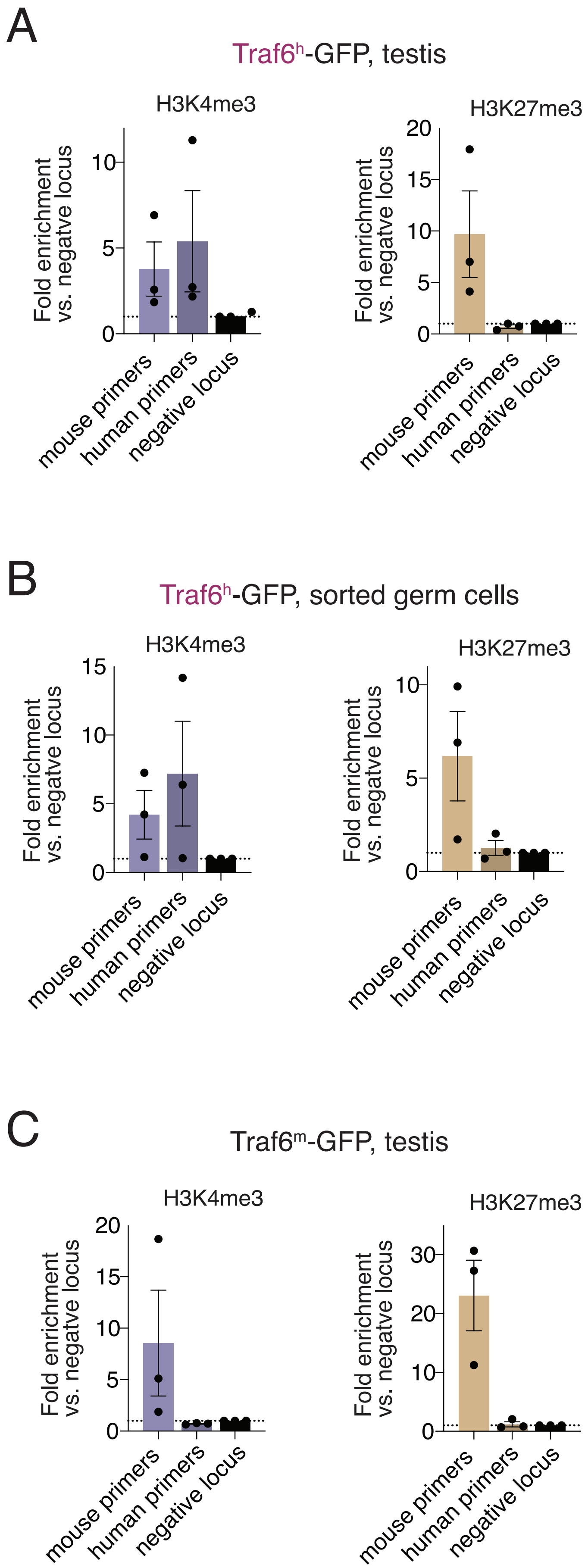

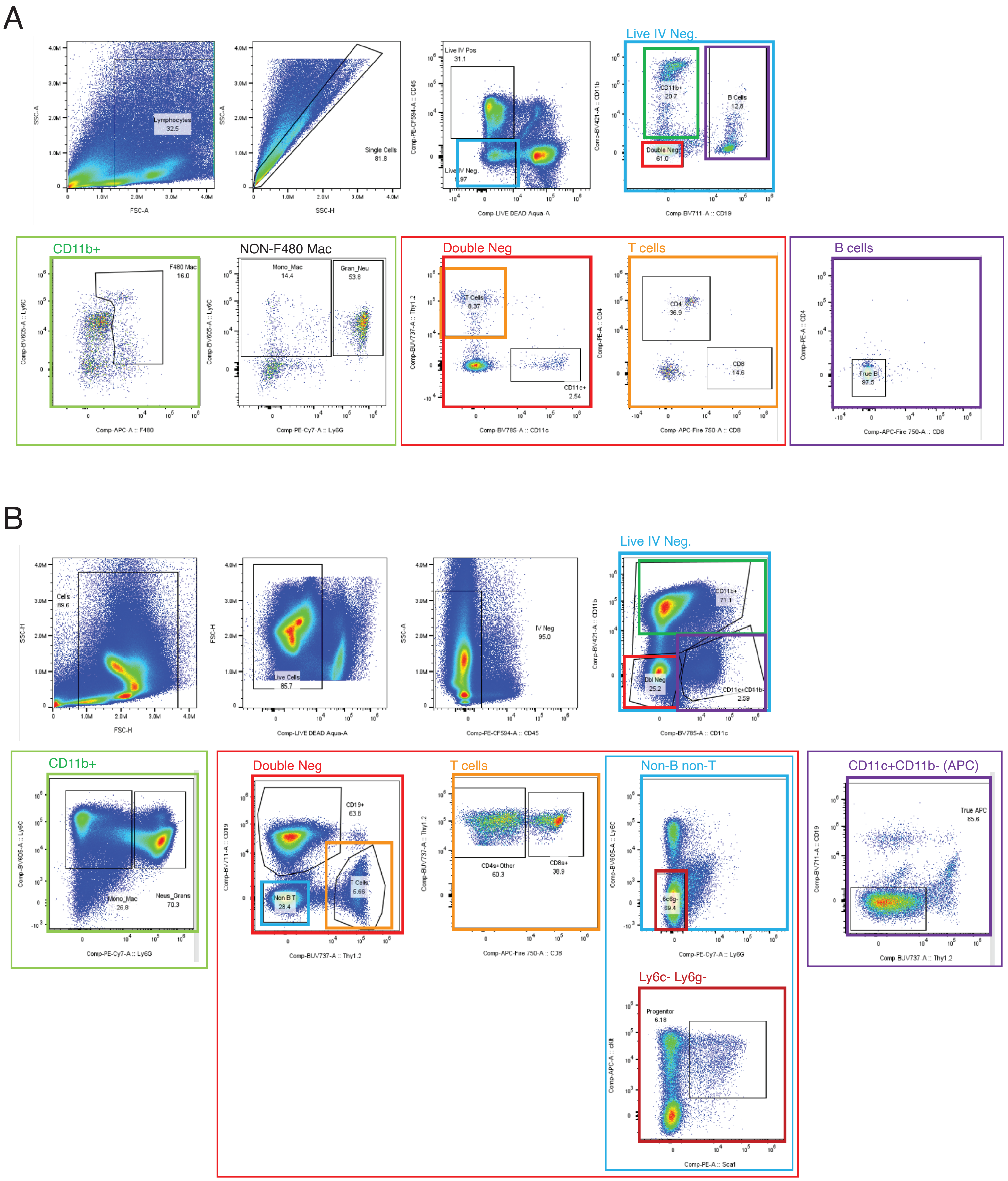

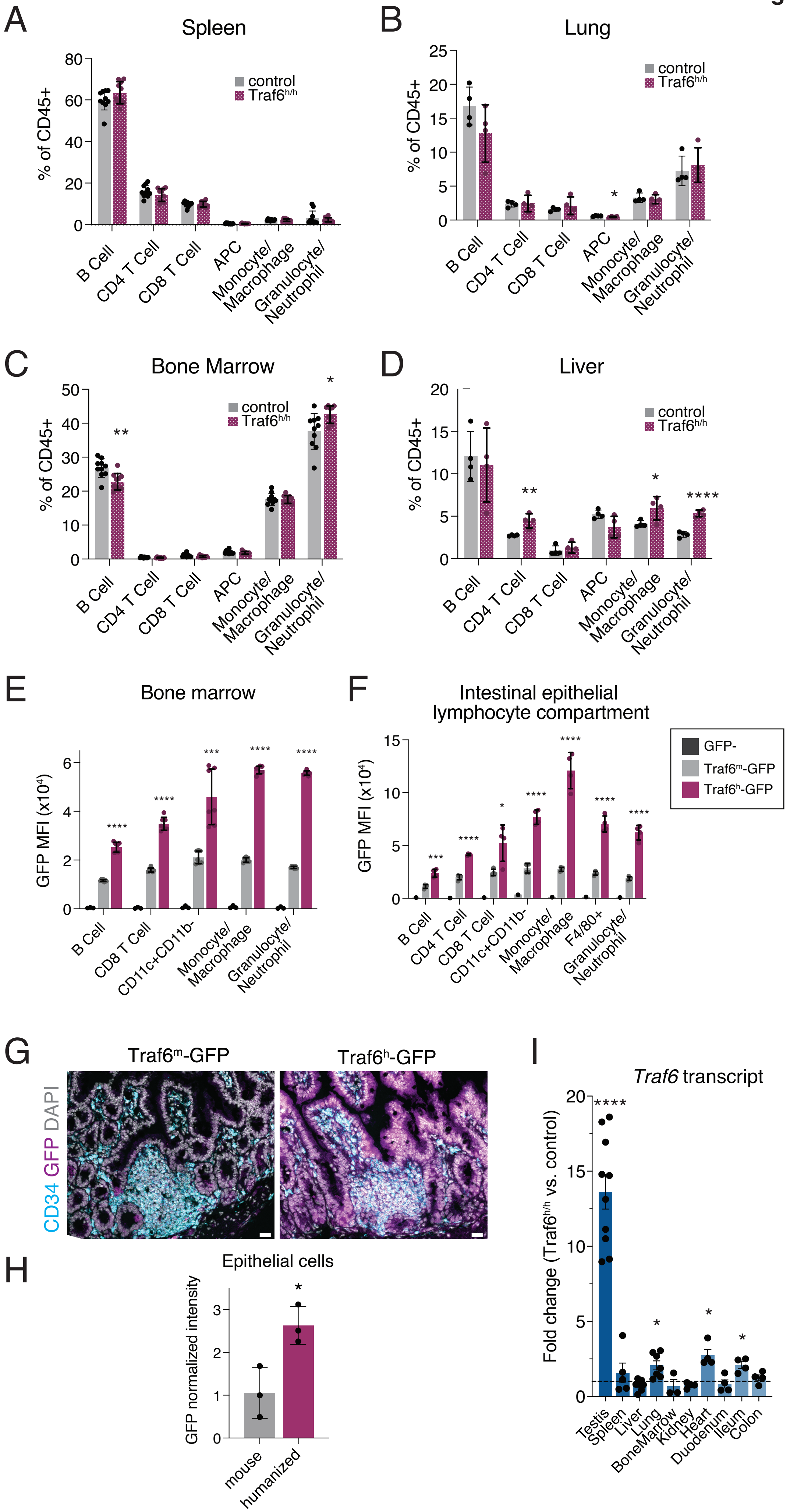

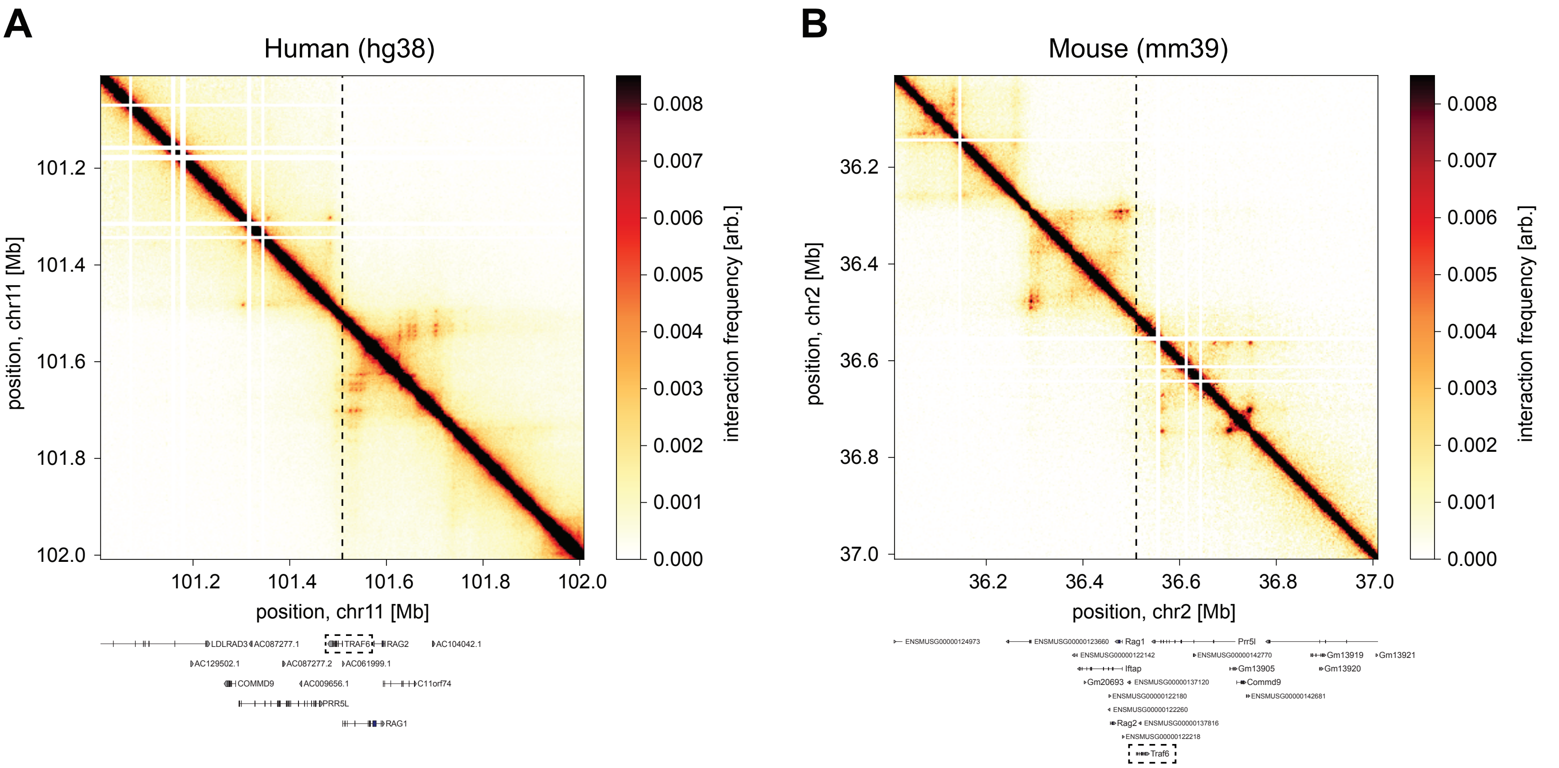

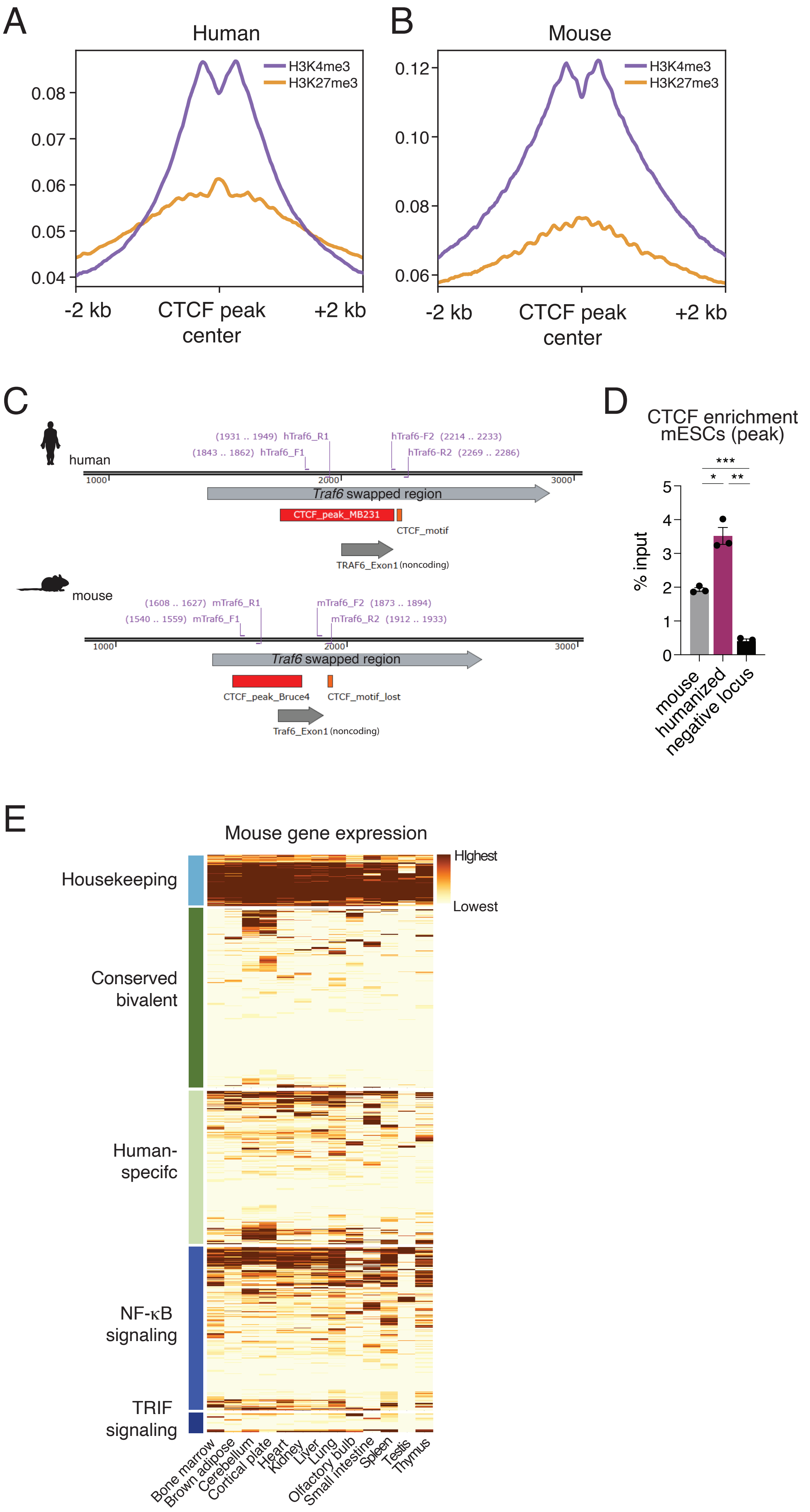
